# Dysregulated mechanoregulation of load-induced bone adaptation in prematurely aged mice: insights from a micro-multiphysics agent-based model

**DOI:** 10.64898/2026.10.22.760951

**Authors:** Daniele Boaretti, Francisco C. Marques, Gisela A Kuhn, Amit Singh, Friederike A. Schulte, Ralph Müller

**Affiliations:** Institute for Biomechanics, ETH Zurich, Zurich, Switzerland

**Keywords:** mechanical loading, PolgA, micro-multiphysics agent-based model, aging, micro-computed tomography

## Abstract

Bone exhibits remarkable ability to adapt to varying mechanical demands. Aging can lead to changes in distribution of cells and proteins that directly affect bone formation and resorption. Given the inherent challenges of quantifying these cellular and protein alterations *in vivo*, innovative *in silico* tools have proven useful to contribute to studying the impact of aging on bone mechanobiology and associated cellular mechanisms. Here, we employed an agent-based *in silico* analysis of *in vivo* micro-computed tomography data of prematurely aged mice to examine the effects of aging and mechanical loading on bone cells and their mechanome, the intrinsic cellular system sensing mechanical forces. Our results from applying a micro-multiphysics agent-based model to an *in vivo* model of premature aging (PolG) showed that aging disrupts mechanoregulation of load-induced bone formation and resorption by eliminating the increase in bone formation through diminished osteoblast activity and mineralization, alongside mechanomic alterations. Specifically, prematurely aged mice exhibited a higher frequency of quiescence at higher mechanical signals and the simulations showed that the number of lining cells and osteoclasts per bone surface increased with age. Normalized trabecular thickness was more positively associated with the ratio of osteoblast to osteoclast number (Ob.N/Oc.N) in the aged control and aged loading groups (R^2^=0.58 and 0.61), than in young control and loading groups (R^2^=0.06 and 0.27), indicating dysregulated remodeling dynamics during aging. In conclusion, our results show that *in silico* tools can give valuable insights into the changes in mechanoregulation and cellular mechanisms occurring under physiological and aging conditions.

## 1 Introduction

Bone remodeling is regulated by bone-forming osteoblasts and bone-resorbing osteoclasts, which maintain the integrity of the skeleton. These cells are also influenced by external factors such as mechanical loading or disease states. While supraphysiological mechanical loading can induce anabolic responses and stimulate bone adaptation (Buenzli et al., 2012; Gould et al., 2021; Lawson et al., 2022), diseases are accompanied by impairments that can also emerge with age, culminating in bone loss and increased fracture risk (Borgstrom et al., 2020). In this regard, investigations of bone cells under such conditions are commonly carried out using *in vivo* experiments (Groessnerschreiber et al., 1992; Schulte et al., 2011; Xiong et al., 2015; McKenzie et al., 2019; Cawley et al., 2020). Critically, in recent decades, computational technologies have surfaced to assist the analysis of such experimental data from a mechanobiological perspective describing bone mechanoregulation. Using micro-finite element analysis (micro-FE), mechanical signals can be computed at the tissue level and correlated with remodeling events determined from time-lapsed *in vivo* micro-CT images (Schulte et al., 2013a; Webster et al., 2015; Scheuren et al., 2020b; Marques et al., 2023). These studies showed that osteoblasts are more likely to form new bone where local mechanical signals are high, while osteoclasts are more likely to resorb bone where mechanical signals are low (Schulte et al., 2013a).

Much of the evidence supporting these principles has been obtained from *in vivo* studies, particularly using genetically modified and naturally aging mouse models, which provide valuable systems for investigating how aging or disease influences skeletal mechanoadaptation. Among these, the PolgA^(D257A/D257A)^ model, a mouse model of premature aging associated with mitochondrial dysfunction, is characterized by accelerated accumulation of mitochondrial DNA mutations and the development of multiple age-associated phenotypes. Although this model does not recapitulate all aspects of chronological aging, it provides a mechanistically defined model to investigate the contribution of mitochondrial dysfunction to age-related changes in bone mechanobiology. Previously, we showed that in PolgA^(D257A/D257A)^ mice, mechanical loading alone was insufficient to stimulate load-induced bone formation (Scheuren et al., 2020a). Even in wild type (WT) mice, if the frequency or magnitude of the applied load was too low, bone formation was only weakly promoted (Scheuren et al., 2020b). Therefore, during aging, factors beyond the mechanical stimulus itself may impact the biochemical environment and single-cell mechanomic processes underlying the translation of mechanical signals into bone remodeling responses. To advance this exploratory analysis of bone mechanobiology, micro-multiphysics agent-based (micro-MPA) *in silico* modeling (Borgiani et al., 2017; Lafuente-Gracia et al., 2021; Tourolle et al., 2021; Boaretti et al., 2022; Ledoux et al., 2022) has emerged as a powerful and versatile tool. Here, we investigated how aging alters the cellular and mechanical mechanisms underlying bone remodeling and mechanoregulation, using a recently established micro-MPA *in silico* model (Boaretti et al., 2023). We used time-lapsed *in vivo* micro-CT images of the sixth caudal vertebra (CV6) of PolgA^(D257A/D257A)^ to recreate the factorial experimental study using in a previous *in vivo* study in which supraphysiological mechanical loading and aging were the experimental factors (Scheuren et al., 2020a). We hypothesized that aging diminishes cellular contributions to remodeling, including osteoblast activity, mineralization, and the cell mechanome, thereby changing the relationship between mechanical signals and formation or resorption events, leading to a diminished bone response to mechanical loading. To test this hypothesis, we evaluated *in vivo* bone mechanoregulation during aging and compared these responses with those predicted *in silico*.

Our results obtained from *in silico* data showed that disrupted mechanoregulation of bone formation and resorption during aging was characterized by reduced osteoblast activity and mineralization, as well as altered cellular and mechanomic patterns of bone remodeling meaning that quiescent bone surfaces were more frequently associated with higher mechanical signals, and lining cells and osteoclasts per bone surface were increased. We conclude that computational analysis of *in vivo* bone dynamics can provide valuable complementary mechanistic insights into the cellular and mechanical regulation of bone adaptation.

## 2 Methods

### 2.1 *In vivo* mouse data

The original *in vivo* data employed in this analysis were collected and published previously (Scheuren et al., 2020a). Briefly, the mice were grouped based on their age (young mice: 15 weeks old, aged mice: 38 weeks old). Mechanical loads were applied to the sixth caudal vertebrae of each mouse (CV6) via stainless steel pins (Fine Science Tools, Heidelberg, Germany) inserted in the fifth and seventh vertebrae following the protocol previously established (Webster et al., 2008). Either sham (0N control) or 8N cyclic (10 Hz) loading was applied to the sixth vertebra for five minutes, three times per week over 4 weeks. Weekly *in vivo* micro-CT images (vivaCT40 or vivaCT80, Scanco Medical AG, Brüttisellen, Switzerland) were acquired with an isotropic voxel resolution of 10.5 µm. Eventually, five time-lapsed images of each vertebra were obtained for the four groups of animals: young loading, young control, aged loading, and aged control mice. All mouse experiments reported in the present study were previously carried out in strict accordance with the recommendations and regulations in the Animal Welfare Ordinance (TSchV 455.1) of the Swiss Federal Food Safety and Veterinary Office.

### 2.2 *In silico* micro-MPA model

In this study, we simulated trabecular bone remodeling using a micro-MPA *in silico* model that was previously established for simulating homeostatic trabecular bone remodeling (Boaretti et al., 2023). The aged groups had a higher average trabecular bone volume fraction (BV/TV) than the young groups *in vivo* at the beginning of the study (23% and 16% for aged and young mice, respectively). Therefore, considering such an increase in the BV/TV during natural growth, we chose to have the same initial bone microarchitecture in a simulation of control or loading for a given age. Thus, from each group of control animals, we extracted five mice (n=5) who had the minimum distance from the median of the rescaled BV/TV to their initial value (Norm. BV/TV) of the group over the four weeks. Then, at a given age, we used this subgroup from the control group as input to simulate both the control and loading groups. For each of the four groups of *in vivo* data, a configuration of the model was defined and used for simulating bone remodeling for four weeks, which corresponds to the time frame of the *in vivo* study. Each configuration required an initial pre-run of the model using the parameters from the configuration of the control group at the corresponding age of the input data to let the model enter an active state. We assumed the response to mechanical loading would have been the same between the mice belonging to the control and loading groups, if they received the same loading, and we tested the effect of mechanical loading on the same mouse that belonged to the control group *in vivo*. The description of the micro-FE analysis, the extension of the model for simulating mechanical loading and aging in PolgA mice, and the configurations of the four branches with their related parameters are described in the Supplementary material in Table S1.

### 2.3 Bone morphometry, cell numbers and cytokine concentrations

We analyzed the *in vivo* data and *in silico* results using static and dynamic bone morphometry. We computed the static parameters for each day of the simulation and for each weekly *in vivo* micro-CT image, the dynamic parameters by using a two-week time period to determine the formed, quiescent, and resorbed surfaces and volumes of trabecular bone. We used the image processing language XIPL (Hildebrand et al., 1999) with the same procedures previously employed in the analysis of the *in vivo* data used as input for this study. We analyzed BV/TV, specific bone surface (BS/BV), trabecular thickness (Tb.Th), trabecular separation (Tb.Sp), and trabecular number (Tb.N). BV/TV, Tb.Th, Tb.Sp, and Tb.N are the normalized values to their initial values for each sample. The dynamic parameters analyzed were bone formation rate per bone volume (BFR), bone resorption rate per bone volume (BRR), mineral apposition rate (MAR), mineral resorption rate (MRR), mineralizing surface per bone surface (MS), and eroded surface per bone surface (ES).

At the end of the simulation, we used a representative sample of each simulated group to extract a cross-section of the trabecular region. We plotted the concentrations of osteoprotegerin (OPG), sclerostin (Scl), and receptor activator of nuclear factor kappa-B ligand (RANKL) and the positions of osteoblasts, osteoclasts, and lining cells with trabecular bone. The cell numbers of osteoblasts, lining cells, and osteoclasts (Ob.N, Li.Ce.N, and Oc.N respectively) were computed for each day of the 4-week simulation for each sample in every simulated group. Cell numbers were determined by dividing the total number of cells in the trabecular region of the simulation by the bone surface at the corresponding time point. Similarly, the spatial average concentration of OPG, Scl, and RANKL was computed in the whole trabecular region for each day of the simulation and for each sample of the four groups of simulations. Univariate linear regression was used to predict BV/TV, Tb.Th, Tb.Sp, Tb.N, and BS/BV with cell numbers and cytokine concentrations.

### 2.4 Mechanoregulation analysis

The mechanoregulation analysis of the *in silico* and *in vivo* data followed the same approach used in a previous study (Schulte et al., 2013a) and it used the same remodeling regions defined above. Given a baseline micro-CT image with its micro-FE results and the subsequent weekly micro-CT image, the micro-FE results and remodeling regions were used again for determining the conditional probability of observing bone formation, resorption, and quiescence on the surface based on the local mechanical signal. Strain energy density in bone (SED_b_) and the norm of the gradient of strain energy density in the marrow (∇SED_m_) were used as mechanical signal in this analysis as performed in a previous study (Marques et al., 2023). The analysis of SED_b_ and effective strain (EFF_b_) required using the voxels on the bone side of the surface, whereas the analysis of ∇SED_m_ required using the voxels on the marrow side of the surface. The conditional probabilities were defined for every 1% binned value of the rescaled mechanical signal, SED_b_/SED_bmax_, EFF_b_/EFF_bmax_, and ∇SED_m_/∇SED_mmax_, where SED_bmax_, EFF_bmax_, and ∇SED_mmax_ were the 99th percentile of the SED_b_, EFF_b_, and ∇SED_m_ in the micro-FE results of the baseline image of this comparison. This procedure was applied for the first weekly comparison and analyzed collectively for each group of mice using averages of each conditional probability over the relative rescaled range of the mechanical signal.

The first two time points of the *in vivo* data were further analyzed. In this restricted period for each group, we computed the conditional probability of the net response by subtracting the conditional probability of resorption from the corresponding conditional probability of formation for each binned value of the rescaled mechanical signal. Then, the conditional probabilities of formation, quiescence, and net response were fitted with non-linear regression analysis, hypothesizing an exponential relationship between the rescaled mechanical signal and the conditional probability. With the estimation of the fits, three fitting parameters were obtained for each curve: y_0_, a and b. The meaning of the parameters can be directly linked to the functioning of the mechanosensory system (Schulte et al., 2013a): y_0_, a, and b represented the probability of non-targeted remodeling, the mechanical sensitivity, and the mechanical control in remodeling, respectively. y_0_ represents level of remodeling in the absence of mechanical stimulus (i.e., at SED=0). Coefficient a represents the magnitude of the response, i.e., the difference between the initial and maximum formation probability or the initial and minimum resorption probability. Coefficient b represents the rate of change, indicating how quickly the formation probability increases or the resorption probability decreases as SED increases. The R^2^-values were computed to assess the quality of the fits.

### 2.5 Receiver operator characteristic analysis

The remodeling regions of formation and resorption from the *in vivo* data were obtained by registration and overlay of subsequent weekly time-lapsed images. The remodeling images from the *in silico* data were already available from the bone morphometry analysis we described above. The micro-FE analysis was computed for each baseline image of the weekly comparison for the *in vivo* data (Supplementary Material), while this information was retrieved from the output of the micro-MPA simulations. The trabecular region was extracted for each sample using masks obtained with an established procedure from every weekly micro-CT image until the second to last one (Lambers et al., 2011). The trabecular region was used to extract the remodeling information and the SED values from the whole image of each vertebra.

We used a receiver operator characteristic (ROC) analysis to determine the predictive performance of SED and ∇SED for bone formation and bone resorption events *in vivo* in the trabecular region between subsequent time points, adopting the approach used in a previous work (Scheuren et al., 2020b). Briefly, we ran this evaluation on all the pairs of images between consecutive time points. This is a method to detect the performance of a binary classifier of an event. This analysis was carried out on the trabecular surface regions, analyzing the performance of SED and ∇SED as predictors. For SED, we used the voxels on the bone side of the bone-marrow interface (SED_b_), whereas we used the voxels on the marrow side of the interface between bone and marrow for ∇SED (∇SED_m_). For every threshold of the classifier, a true positive rate and a false positive rate were computed (Fig. S1). By varying the threshold of the classifier from its minimum to its maximum value, the ROC curve plots the true positive rates versus the false positive rates. Using this approach, we computed the area under the curve (AUC). The AUC provides a measure of the model’s ability to discriminate between two outcomes. An AUC of 0.5 corresponds to discrimination at random chance, whereas an AUC of 1.0 indicates perfect discrimination.

Additionally, along with the AUC, we computed the optimal threshold of formation and resorption by maximizing the differences between the true-positive and false-positive rates. This optimization corresponded to the point with the maximum distance from the ROC curve of a random classifier, i.e., maximizing the difference between the true positive and false positive rates. After computing the optimal threshold for a predictor of bone formation (OTF), the voxels with a mechanical signal above this threshold were considered formed according to that predictor. On the other hand, after computing the optimal threshold for a predictor of bone resorption, the voxels with a mechanical signal below this threshold (OTR) were considered to be resorbed according to that predictor. For a given sample, each threshold was rescaled to the threshold of the first weekly comparison to define the normalized optimal threshold for formation and resorption (Norm.OTF and Norm.OTR, respectively).

### 2.6 Statistical Analysis

For the ROC and mechanoregulation analysis of the mechanical signals, we used Python 3 (Python Software Foundation, www.python.org). The source code of the micro-MPA model was developed in C++ and wrapped in Python, in line with the previous version of the model used for simulating homeostasis (Boaretti et al., 2023). Similarly, the computational platform used for running the simulations was the Swiss National Supercomputing Center (CSCS, Lugano, Switzerland). A complete simulation required 3 to 5 hours, taking advantage of the parallel computing capabilities through 8 nodes (Piz Daint, a Cray XC40/50 model) for the simulation and 2 nodes for the micro-FE analysis of the vertebra.

We used R (R Core Team (2019), R Foundation for Statistical Computing, Vienna, Austria) for the statistical tests of the AUC, Norm.OTF, Norm. OTR, the static and dynamic bone morphometric parameters obtained from the *in vivo* and *in silico* data. Repeated measures ANOVA implemented as a linear mixed model with the lmerTEST package (Kuznetsova et al., 2017) was used. With this test, we detected if significant differences were present due to the effect of time, group (*in silico* vs. *in vivo* or Norm.OTF vs. Norm. OTR), and the interaction between time and group, which indicated whether the variable developed differently over time between groups. Pairwise t-test followed by Cohen’s D were computed to determine the standardized difference between the AUC computed with SED_b_ and ∇SED_m_. Additionally, we computed the p-value for each univariate linear regression to determine if the slope was significantly different from zero. P-values smaller than 0.05 were considered significant. The mean and the standard deviation of the selected data, or alternatively boxplots, are plotted in the figures. Visualizations of the trabecular 3D bone microarchitecture were generated using ParaView (Kitware, Version 5.10; Clifton Park, NY).

## 3 Results

### 3.1 Age-related changes in bone morphometry

Bone morphometry was first assessed in the generated structures (Fig. 1A). Normalized BV/TV showed no significant differences between *in silico* and *in vivo* data (Fig. 1B). Tb.Th was generally consistent across groups and time points, except for a significant temporal change in the young loading group (Tables S3–S4). In aged mice, remodeling did not significantly alter BV/TV or Tb.Th (Fig. 1B and Fig. S2), while Tb.N was overestimated and BS/BV and Tb.Sp were underestimated relative to *in vivo* measurements (p < 0.05).

**Figure 1.**
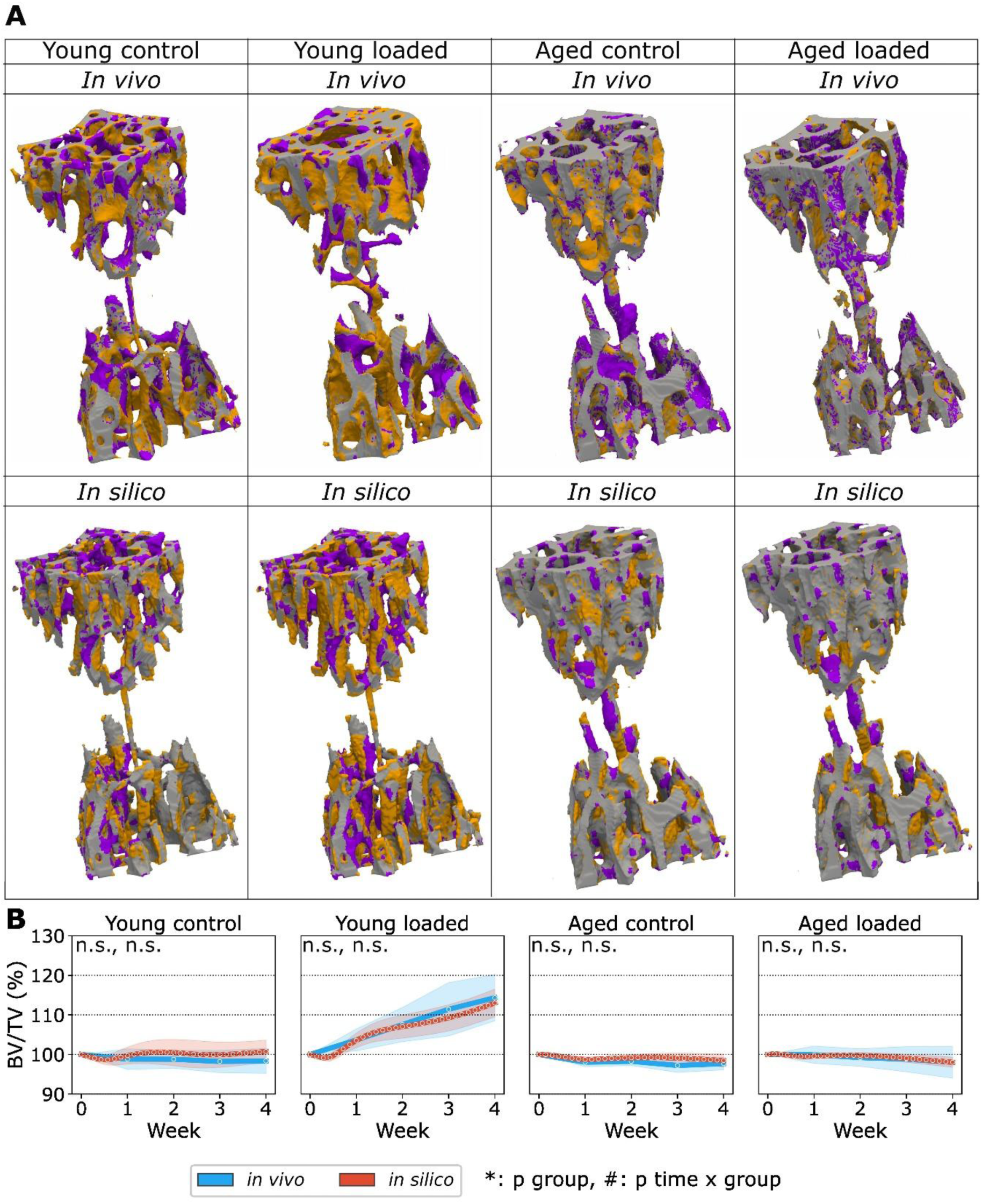
*In vivo* and *in silico* trabecular bone remodeling in the four groups of mice. (A) Representative overlay of thresholded micro-computed tomography images from the four groups of mice showing sites of formation (orange), quiescence (grey), and resorption (violet) in representative *in silico* and *in vivo* data between week 0 and 4. The first row shows the remodeling sites of *in vivo* samples, the second row from the simulations. (B) Normalized trabecular bone volume fraction (Norm. BV/TV) in the four groups of mice *in silico* and *in vivo* over four weeks. The *in vivo* curves show the data from the time-lapsed weekly images. The *in silico* data are reported on a daily basis. Light blue = *in vivo*, Red = *in silico*. n=5 *in silico* and *in vivo* control groups, n=9 *in vivo* loaded groups. (n.s., * p<0.05, ** p<0.01, *** p<0.001, **** p<0.0001 for the group comparison, # p<0.05, ## p<0.01, ### p<0.001, #### p<0.0001 for the interaction time*group determined by two-way ANOVA).

The agreement in dynamic remodeling indices varied with age and loading condition (Tables S1–S4). In young control mice, the model reproduced BFR and MAR, as well as the temporal trends in MS and ES, although MS and ES were underestimated on average. BRR and MRR showed partial agreement with experimental data. In young loaded mice, BFR, BRR, and ES agreed with *in vivo* data, while the temporal trends in MAR and MS were captured despite differences in their average values. Conversely, MRR showed agreement in average values but not in its temporal trend. In aged control mice, BRR was the only dynamic index without significant differences from *in vivo* data. In aged loaded mice, BRR showed no significant differences in either average values or temporal evolution. Overall, the model reproduced normalized bone volume and selected remodeling dynamics, but discrepancies in trabecular architecture and the age-dependent agreement of dynamic indices highlight limitations in capturing bone remodeling across physiological conditions.

### 3.2 Cellular behavior in prematurely aged mice compared to young mice

We analyzed the cellular environment of osteoblast, osteoclast, and lining cells in the trabecular region (Fig. 2). The young control group displays the number of osteoblasts in homeostatic conditions, while mechanical loading increases the number of osteoblasts through enhanced cellular proliferation and activity (Fig. 2A). The aged control and loading groups exhibited similar and slightly lower cell populations, with more lining cells under loading than in the control condition. The average osteoblast number per bone surface (Ob.N/BS) showed small fluctuations around the initial value in the simulations of the young control group, and considerable increased in the young loading group (Fig. 2B). Conversely, the simulations of the aged groups showed a reduction in this number. The average lining cell number per bone surface (Li.Ce.N/BS) remained very low in the simulations of the young control group, while it showed an increase over time in the simulations of the young loading group (Fig. 2B). The simulations showed that the number of lining cells per bone surface increased with age, with an initial increase followed by a decrease. The average osteoclast number per bone surface (Oc.N/BS) fluctuated over time in the simulations of the young control group, and in the young loading group it increased by 2-fold (Fig. 2B). The aged groups showed a higher Oc.N/BS from the beginning, with an increasing trend over time.

**Figure 2.**
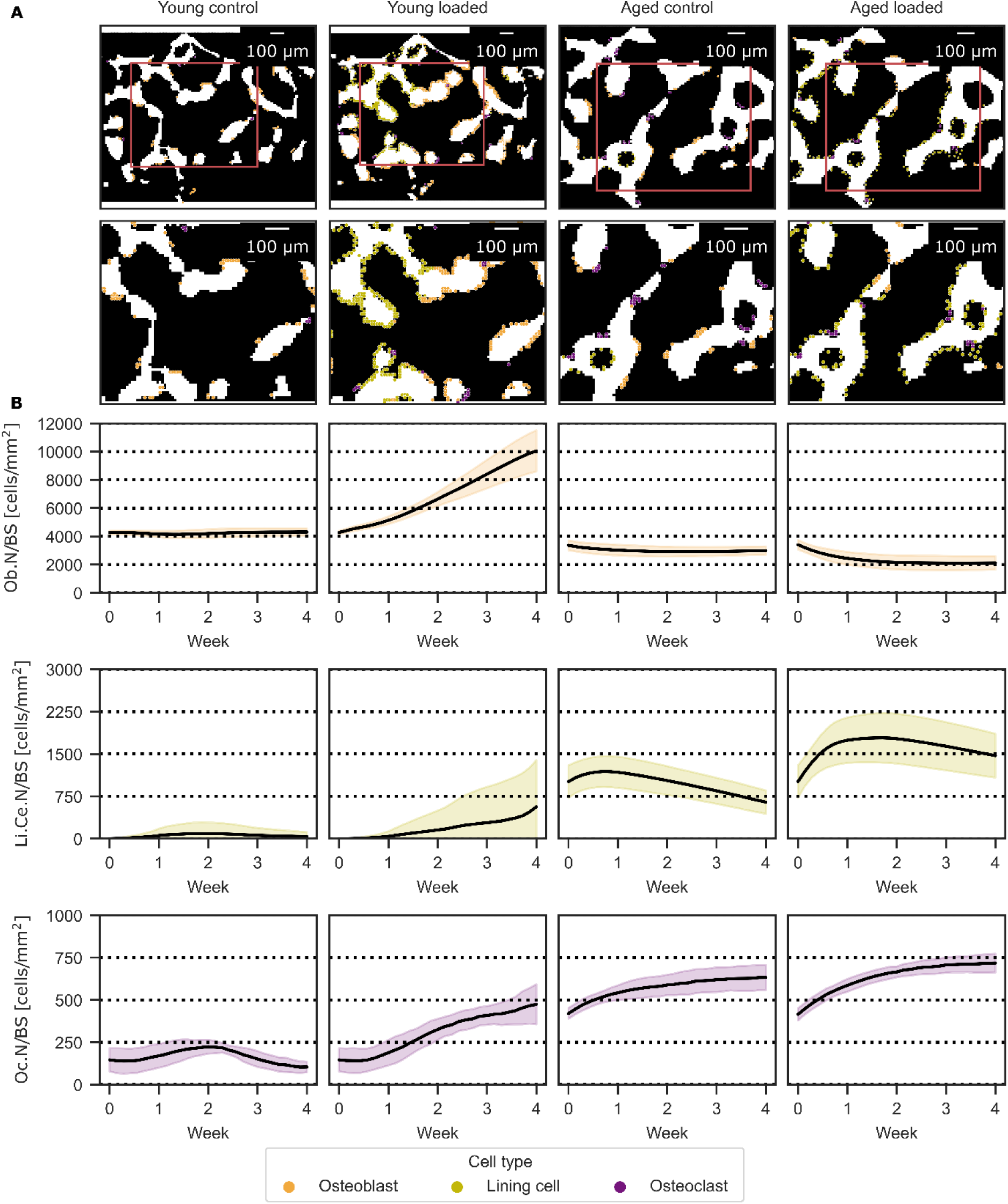
Cell distributions from the micro-MPA simulations. (A) Cross-section of the trabecular region showing osteoblasts (orange), lining cells (olive yellow), and osteoclasts (purple) at the final time point of the simulation from a representative sample of each simulated group. (B) Averages and standard deviations of osteoblast, lining cell, and osteoclast cell numbers per bone surface (Ob.N/BS, Li.Ce.N/BS, and Oc.N/BS) in the micro-MPA simulations of the four groups.

### 3.3 Predicted serum cytokine concentrations

OPG, Scl, and RANKL concentrations computed from the micro-MPA simulations are shown in Fig. 3 with the same representative samples and time point as in Fig. 2 being used. For visualization purposes, only the bone pixels show the concentration value; nonetheless, the concentrations were computed everywhere in the simulation domain of the micro-MPA model. The young loading group exhibited the highest OPG values compared to the other groups due to higher load-induced production of OPG (Fig. 3A). The concentration values were higher in the aged loading group than in the aged control group. The Scl concentration was relatively low in the young control group compared to the young loading group (Fig. 3A). Similarly high values were also obtained in both aged groups. Likewise, the young control group showed a lower RANKL concentration, and the young loading group higher values (Fig. 3A). When compared to the aged loading group, the aged control group had a higher concentration of RANKL. The average trabecular concentration of OPG, Scl, and RANKL was computed over time for each day of the simulation and for each sample (Fig. 3B). Here, the young groups showed a decrease over time of the initial concentration, while the aged control group had a relatively constant concentration over time. OPG concentration increased in the aged loading group. Scl concentration increased over time for both the young control and loading groups, and remained relatively constant over time in the aged groups. Additionally, RANKL concentration had a little fluctuation over time in the young control group, while with loading it increased. In the aged groups, it had a higher initial value and decreased over time.

**Figure 3.**
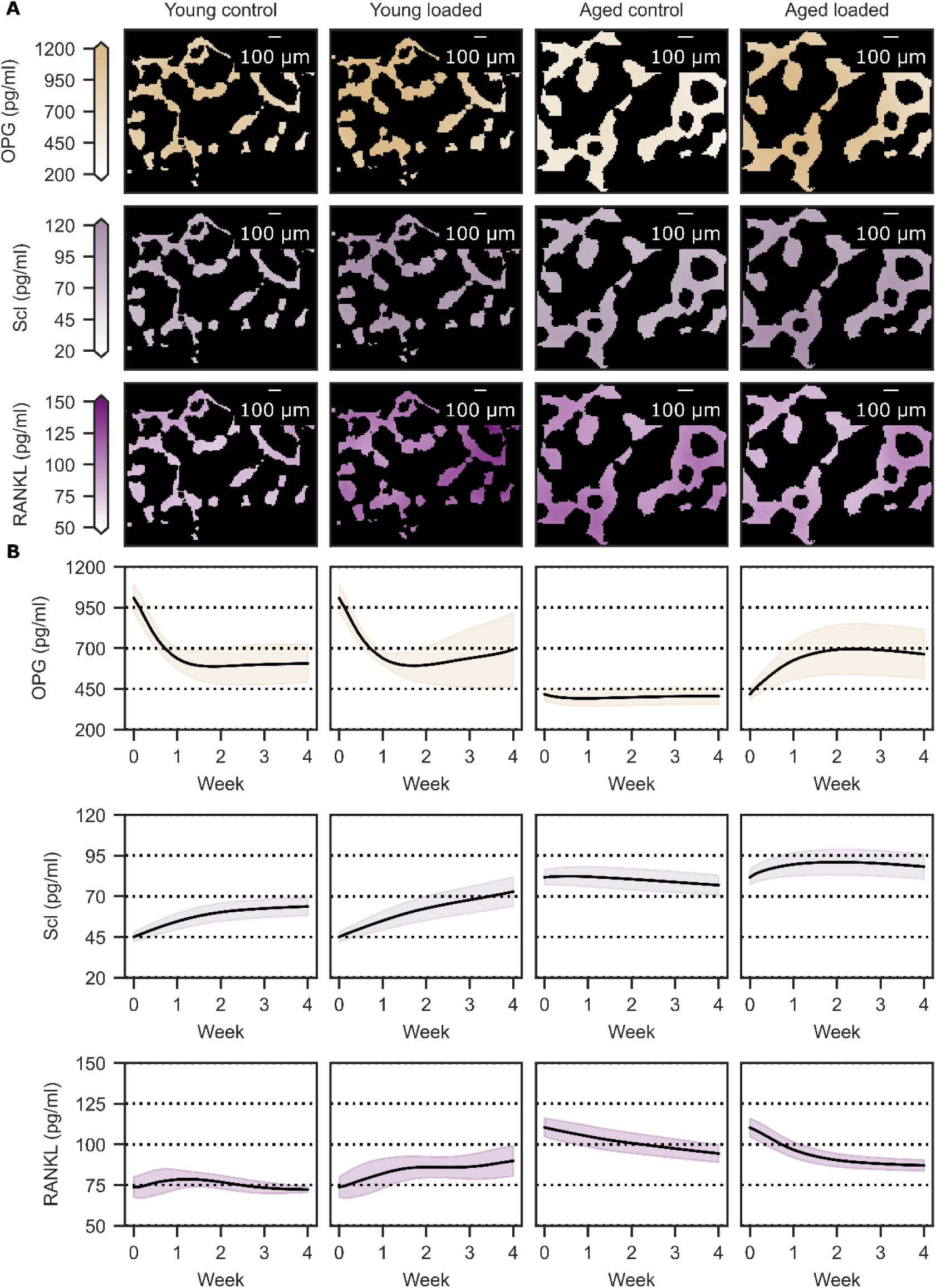
Cytokine distributions from the micro-MPA simulations. (A) OPG, Scl, and RANKL concentration in bone at the final time point of the simulation from a cross-section of the trabecular region collected from a representative sample of each simulated group. (B) Averages of OPG, Scl, and RANKL concentrations in the micro-MPA simulations of the four groups.

### 3.4 Regression analysis

The most relevant findings from the univariate linear regression analyses were obtained when predicting Norm. Tb.Th from the osteoblast-to-osteoclast number ratio (Ob.N/Oc.N), osteoclast number per bone surface (Oc.N/BS), and the average sclerostin (Scl) concentration in the trabecular region (Fig. 4 and Table S5). Norm. Tb.Th was positively associated with Ob.N/Oc.N in the aged control and aged loading groups, with the regression explaining 58% and 61% of the variability, respectively (R^2^=0.58 and 0.61). In contrast, Ob.N/Oc.N explained little variability in the young control group (R^2^=0.06), while a positive association was observed in the young loading group (R^2^=0.27). The regression slopes were negative in the remaining groups. Oc.N/BS showed a strong positive association with Norm. Tb.Th in the young loading group (R2=0.81), whereas little variability was explained in the young control group (R^2^=0.04). In the aged control and aged loading groups, the association was negative, with R^2^=0.54 and 0.57, respectively. Finally, Scl showed a positive association with Norm. Tb.Th only in the young loading group (R^2^=0.51). The relationship was weaker in the young control group (R2=0.23) and negligible in the aged control and aged loading groups (R^2^=0.01 and 0.18, respectively). Notably, the regression slopes for the aged groups were zero or negative, whereas both young groups showed positive slopes. Overall, these findings suggest that Ob.N/Oc.N, Oc.N/BS, and Scl are potential markers for characterizing loading-induced and age-related alterations in trabecular bone remodeling identified by the micro-MPA simulations.

**Figure 4.**
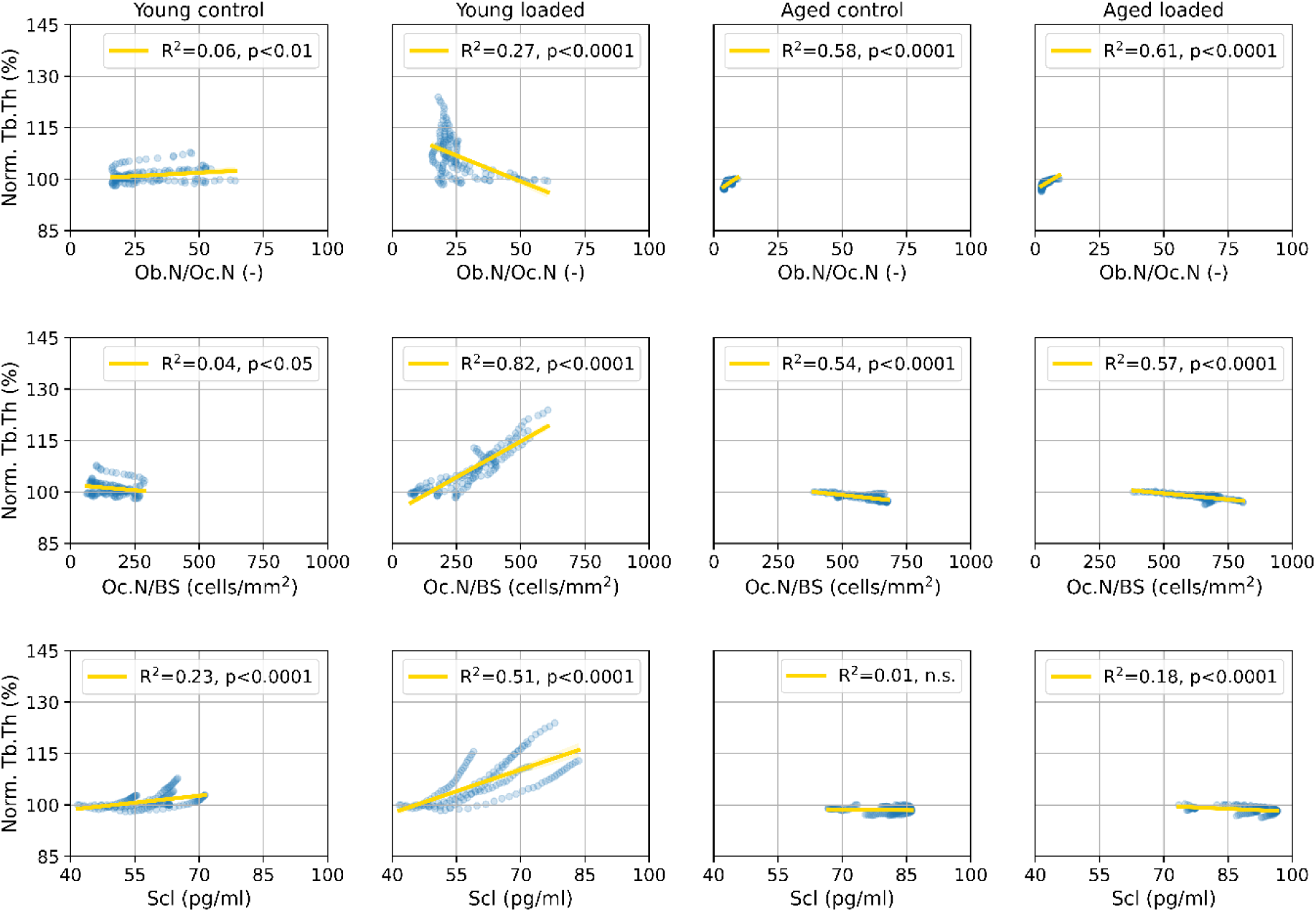
Analysis between trabecular thickness and markers from micro-MPA simulations. Linear regression of selected output variables (Ob.N/Oc.N, Oc.N/BS, and Scl) against the normalized trabecular thickness (Norm. Tb.Th) from the micro-MPA simulations of the four groups of mice. A single dot represents the values for a single sample from a single day during a simulation of four weeks.

### 3.5 Aging disrupts the anabolic response to mechanical loading and mechanoregulation in prematurely aged mice

Across all groups, SED_m_ best predicted bone formation AUC= 0.62, while SED_b_ best predicted resorption (AUC= 0.61; Fig. 5A and Table S6). Optimal formation (Norm.OTF) and resorption (Norm.OTR) thresholds remained constant over time in control and aged mice (Tables S7 and S8). However, mechanical loading in young mice significantly reduced both thresholds over time (p < 0.0001; Table S6), indicating an adaptive response that was entirely absent in aged loaded mice.

**Figure 4.**
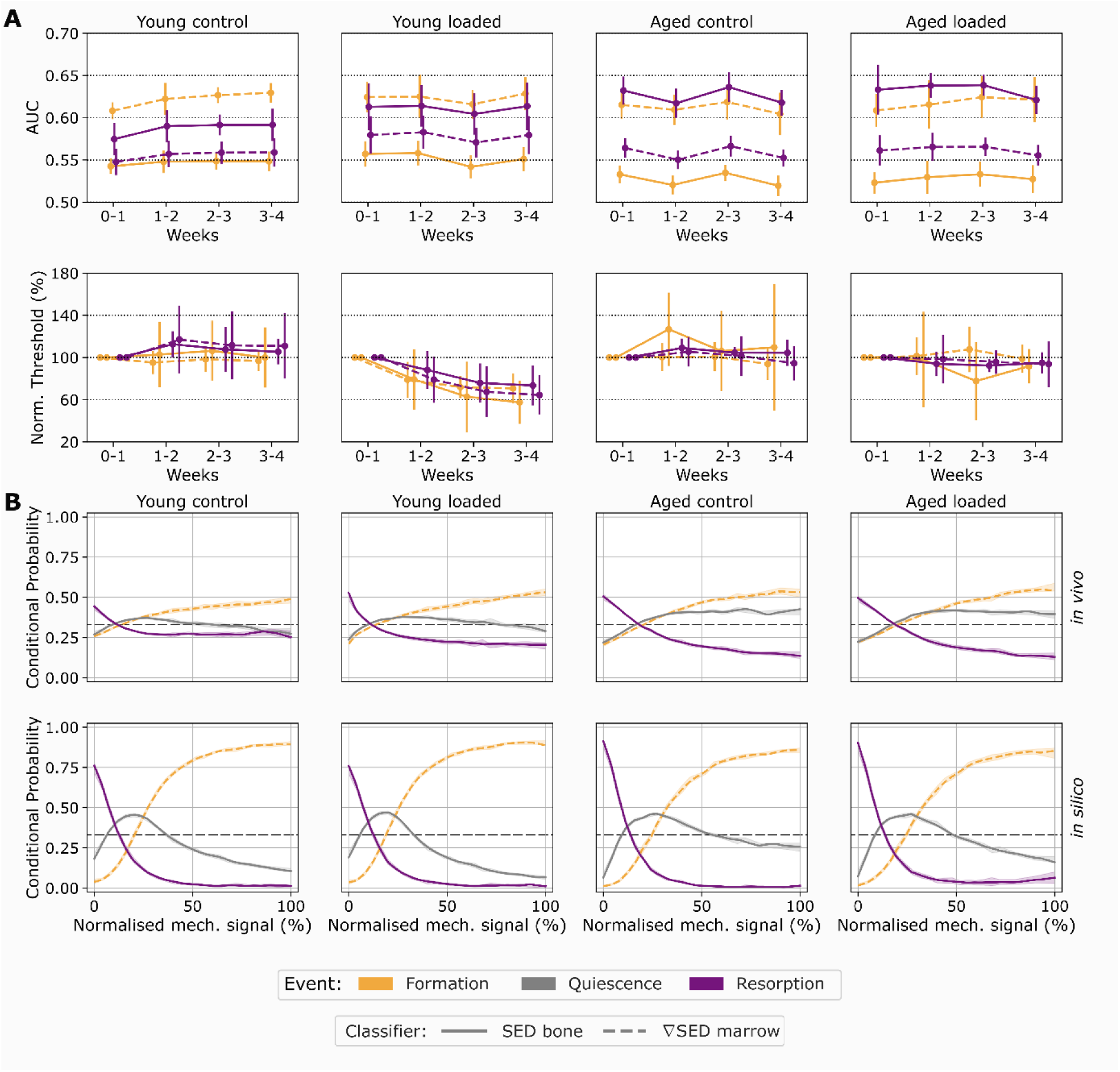
Analysis of trabecular bone remodeling and mechanical environment. (A) The ROC analysis in the trabecular region of the sixth caudal vertebra for the four groups of PolgA mice. The first row shows the area under the curves (AUCs) using strain energy density values in bone (SED_b_) and the norm of the gradient of strain energy density (∇SED_m_) as predictors of the weekly bone formation and bone resorption events *in vivo*. The second row shows the normalized optimal thresholds for bone formation (OTF) and bone resorption (OTR) using SED_b_ over time. (B) Mechanoregulation analysis in the trabecular region of the sixth caudal vertebra for the four groups of PolgA mice *in vivo* (first row) and *in silico* (second row). The mechanoregulation curves of formation using (∇SED_m_) and resorption and quiescence using SED_b_ for the first weekly interval are shown. The dashed horizontal black line represents the random probability of observing one of these three events, which corresponds to 0.33.

*In vivo*, mechanical loading in young mice increased formation and quiescence probabilities while decreasing resorption (Fig. 5B). Conversely, aged mice showed no loading-induced increase in bone formation, maintaining elevated quiescence across all mechanical signal levels and higher resorption at low signal levels (Fig. 5B and Fig. S3). Micro-MPA simulations replicated these patterns, capturing clear signal-dependent separation between formation and resorption in young mice, but reflecting shifted thresholds and increased quiescence in aged mice (Fig. 5B and Fig. S4).

Exponential curve fitting confirmed that loaded aged mice experienced a 3-fold reduction in mechanical control for bone formation using SED compared to controls (Table S9). Together, these results demonstrate that premature aging is associated with reduced mechanical sensitivity and impaired load-induced bone adaptation.

## 4 Discussion

In this computational study, using a micro-MPA model which includes the actions of the bone cells modeled as agents that can produce cytokines and remodel bone through changes in the mechanome around a single cell, we found that mechanoregulation of bone formation and resorption was altered with aging. Simulations identified reduced osteoblast activity and mineralization in aged mice, which showed higher frequency of quiescence at higher mechanical signals than in young mice. Our simulations reproduced the experimentally observed impaired anabolic response associated with aging, alongside the absence of bone growth response to mechanical loading in aged mice. These results were in strong agreement with our previous experimental findings, reinforcing the robustness and reliability of the agent-based micro-MPA model. Moreover, SED_b_ was a better predictor of bone resorption than ∇SED_m_, while ∇SED_m_ was a better predictor of bone formation than SED_b_ (Fig. 5, Table S6), suggesting that distinct mechanical signals may contribute to the regulation of these two remodeling processes.

The aged phenotype in the simulations was reproduced by reducing cellular activity, especially the osteoblastic cells, resulting in impaired bone formation due to a reduced MS and lower MAR, which are correlated to the mineralization rate r^mineral^, the osteoblast availability, and the synthesis rate of osteoid β_Oc^(mineral,osteoid)^. These changes are consistent with previous observations of reduced MSC and osteoblast proliferation rates, osteoblast activity, and reduced mineralizing surface in aged PolgA mice(Meakin et al., 2014; Gu et al., 2018; Meng et al., 2018; Corrigan et al., 2019; Dobson et al., 2020; Scheuren et al., 2020a). The reduced osteoclastic apoptosis rate in the simulations was also consistent with the lower *in vivo* MRR observed in the aged mice.

The detection of ∇SED_m_ as a better predictor of bone formation events is in agreement with previous findings (Webster et al., 2015), which found a correlation between the magnitudes of dynamic bone morphometry parameters and the ∇SED_m_. That analysis was more spatially averaged compared to the voxel-based analysis employed in this study, and it showed the correlation between the magnitudes, not the separation of formation and resorption events. In a previous study, ∇SED_m_ was a better predictor for both formation and resorption events in the trabecular region of CV6 in 15-week-old female C57BL/6J mice over a period of four weeks (Scheuren et al., 2020b). Our analysis showed that SED_b_ was a better predictor for bone resorption. It might be that osteocytes from mice of different genotypes could employ a different mechanoregulation (van Hove et al., 2009; Hemmatian et al., 2021). From these results, *in silico* simulations of bone remodeling might be improved by the application of two mechanical signals, SED_b_ and ∇SED_m_, instead of a single one, to regulate bone resorption and bone formation. However, this study did not investigate which cell type should be responsible for the mechanome of those signals or how they could convert them into actual cell activity (Matsuzaka et al., 2021).

The mechanoregulation analysis of the *in vivo* data showed that mechanical loading could promote bone formation and reduce bone resorption at a young age, while aging reduced bone formation and increased quiescence. Furthermore, mechanical loading did not change the conditional probabilities of formation and resorption in the aged loading group, despite the applied mechanical loading condition. This means that altered mechanobiology due to aging might benefit from drug treatment since mechanical stimulation can elicit a reduced anabolic response by itself. The micro-MPA simulation showed an increase in the number of lining cells in both aged groups. Furthermore, it demonstrated that the mechanoregulation curves followed a similar pattern to the *in vivo* data, with quiescence increasing in regions of high mechanical signal. Sclerostin antibody, e.g., romosozumab, treatment or intermittent parathyroid hormone treatment, could recover at least the osteoblast number (Dobnig and Turner, 1995; Kim et al., 2012; Ominsky et al., 2014; Ominsky et al., 2015; Kim et al., 2017). This is in line with previous findings, highlighting that mice with Sost deficiency obtained a higher anabolic response to mechanical loading in cortical bone (Pflanz et al., 2017). The impairment in the response to mechanical loading with age might be due to increased osteoblast’s proliferation with higher strains (Meakin et al., 2014).

In the aged loading group, the temporal evolution of the optimal thresholds was similar compared to both the young and aged control groups (Fig. 5A). Beginning in the second week, we observed bone adaptation in the young loading group, leading to similar optimal thresholds of bone formation and resorption for SED_b_ while the optimal thresholds of ∇SED_m_ slightly decreased over time. In consequence, the young mice were more responsive to loading than the aged mice. This result has implications for bone remodeling simulations in which the concept of the mechanostat is used to study aging. Indeed, previous simulations of bone remodeling that used the mechanostat employed absolute values of SED for all the groups of mice they simulated (Schulte et al., 2013b; Levchuk et al., 2014). The simulation of bone remodeling would require a calibration of the mechanostat thresholds for determining when bone is formed or resorbed in the young and aged groups under the same loading condition. Actually, aged mice have more bone due to natural growth (Scheuren et al., 2020a) and therefore a higher capability of reducing the overall SED and strains (Somerville et al., 2004; Razi et al., 2015b). Moreover, with dysregulation associated with aging, these thresholds might become altered.

The mechanome, as well as other model parameters, were adjusted in the aging simulation to keep the number of altered parameters as low as possible, focusing on the effect of loading on the mechanome. Previous research has suggested that mechanomics can become blurry with less specific osteoblast activation or inhibition (Razi et al., 2015a). There, bone formation and resorption were both affected, hence the sensing of mechanical signals by osteocytes, osteoblasts, and osteoclasts might be altered. The cellular activity and signaling pathways might be affected in more aspects than the ones we changed. Indeed, it was observed that the receptor activator of nuclear factor kB ligand-osteoprotegerin (RANKL-OPG) ratio increased with aging, thereby favoring catabolic activity and bone resorption (Chung et al., 2014). Moreover, with aging, the total number of osteocytes was lower, as was the number of sclerostin-positive osteocytes, but it was suggested that an increase in sclerostin synthesis by the osteocytes could compensate for the reduced number of cells (Thompson et al., 2016). On the other hand, Wnts can prevent the apoptosis of osteoblasts and osteocytes (Jilka et al., 2007), meaning that the abundance of sclerostin increases osteoblast and osteocyte apoptosis. More specifically, aging increased cellular apoptosis in PolgA mice (Kujoth et al., 2005). However, the PolgA mouse model does not replicate all cellular aspects of aging, particularly hematopoietic stem cell (HSC) gene expression (Norddahl et al., 2011). Testing all the possible changes in cell activity and signaling pathways would require an extensive simulation set as well as some more specific experimental data to corroborate if studies in different organs or mouse strains are applicable for PolgA mice.

In conclusion, our micro-MPA model and mechanoregulation analysis demonstrated an experimentally confirmed diminished mechanoregulatory response to mechanical loading in prematurely aged mice compared with young mice, with reduced osteoblast activity representing a potential contributing factor. *In silico* tools can complement experimental approaches to improve our understanding of age-related changes in bone mechanobiology while supporting the replacement, reduction, and refinement of animal models.

## 6 Author Contributions

Conceptualization: DB, RM

Methodology: DB, FCM, GAK

Investigation: DB, FCM, GAK

Visualization: DB, FCM, RM

Validation: AS

Supervision: RM, FAS

Writing—original draft: DB

Writing—review & editing: DB, FCM, GAK, FAS, RM

## 8 Acknowledgments

This study was supported by a grant from the Swiss National Supercomputing Centre (CSCS) under project ID s1070.

## References

Boaretti, D., Marques, F.C., Ledoux, C., Singh, A., Kendall, J.J., Wehrle, E., et al. (2023). Trabecular bone remodeling in the aging mouse: A micro-multiphysics agent-based model using single-cell mechanomics. Frontiers in Bioengineering and Biotechnology 11. doi: ARTN 1091294 10.3389/fbioe.2023.1091294.

Boaretti, D., Wehrle, E., Bansod, Y.D., Tourolle Ne Betts, D.C., and Muller, R. (2022). Perspectives on in silico bone mechanobiology: computational modelling of multicellular systems. Eur Cell Mater 44, 56–73. doi: 10.22203/eCM.v044a04.

Borgiani, E., Duda, G.N., and Checa, S. (2017). Multiscale Modeling of Bone Healing: Toward a Systems Biology Approach. Front Physiol 8, 287. doi: 10.3389/fphys.2017.00287.

Borgstrom, F., Karlsson, L., Ortsater, G., Norton, N., Halbout, P., Cooper, C., et al. (2020). Fragility fractures in Europe: burden, management and opportunities. Arch Osteoporos 15(1), 59. doi: 10.1007/s11657-020-0706-y.

Buenzli, P.R., Pivonka, P., Gardiner, B.S., and Smith, D.W. (2012). Modelling the anabolic response of bone using a cell population model. J Theor Biol 307, 42–52. doi: 10.1016/j.jtbi.2012.04.019.

Cawley, K.M., Bustamante-Gomez, N.C., Guha, A.G., MacLeod, R.S., Xiong, J., Gubrij, I., et al. (2020). Local Production of Osteoprotegerin by Osteoblasts Suppresses Bone Resorption. Cell Rep 32(10), 108052. doi: 10.1016/j.celrep.2020.108052.

Chung, P.L., Zhou, S., Eslami, B., Shen, L., LeBoff, M.S., and Glowacki, J. (2014). Effect of age on regulation of human osteoclast differentiation. J Cell Biochem 115(8), 1412–1419. doi: 10.1002/jcb.24792.

Corrigan, M.A., Coyle, S., Eichholz, K.F., Riffault, M., Lenehan, B., and Hoey, D.A. (2019). Aged Osteoporotic Bone Marrow Stromal Cells Demonstrate Defective Recruitment, Mechanosensitivity, and Matrix Deposition. Cells Tissues Organs 207(2), 83–96. doi: 10.1159/000503444.

Dobnig, H., and Turner, R.T. (1995). Evidence That Intermittent Treatment with Parathyroid-Hormone Increases Bone-Formation in Adult-Rats by Activation of Bone Lining Cells. Endocrinology 136(8), 3632–3638. doi: DOI 10.1210/en.136.8.3632.

Dobson, P.F., Dennis, E.P., Hipps, D., Reeve, A., Laude, A., Bradshaw, C., et al. (2020). Mitochondrial dysfunction impairs osteogenesis, increases osteoclast activity, and accelerates age related bone loss. Sci Rep 10(1), 11643. doi: 10.1038/s41598-020-68566-2.

Gould, N.R., Williams, K.M., Joca, H.C., Torre, O.M., Lyons, J.S., Leser, J.M., et al. (2021). Disparate bone anabolic cues activate bone formation by regulating the rapid lysosomal degradation of sclerostin protein. Elife 10. doi: 10.7554/eLife.64393.

Groessnerschreiber, B., Krukowski, M., Lyons, C., and Osdoby, P. (1992). Osteoclast Recruitment in Response to Human Bone-Matrix Is Age-Related. Mechanisms of Ageing and Development 62(2), 143–154. doi: Doi 10.1016/0047-6374(92)90051-E.

Gu, X.H., Fu, X.J., Lu, J.H., Saijilafu, Li, B., Luo, Z.P., et al. (2018). Pharmacological inhibition of S6K1 impairs self-renewal and osteogenic differentiation of bone marrow stromal cells. Journal of Cellular Biochemistry 119(1), 1041–1049. doi: 10.1002/jcb.26272.

Hemmatian, H., Bakker, A.D., Klein-Nulend, J., and van Lenthe, G.H. (2021). Alterations in osteocyte lacunar morphology affect local bone tissue strains. J Mech Behav Biomed Mater 123, 104730. doi: 10.1016/j.jmbbm.2021.104730.

Hildebrand, T., Laib, A., Muller, R., Dequeker, J., and Ruegsegger, P. (1999). Direct three-dimensional morphometric analysis of human cancellous bone: microstructural data from spine, femur, iliac crest, and calcaneus. J Bone Miner Res 14(7), 1167–1174. doi: 10.1359/jbmr.1999.14.7.1167.

Jilka, R.L., Weinstein, R.S., Parfitt, A.M., and Manolagas, S.C. (2007). Quantifying osteoblast and osteocyte apoptosis: Challenges and rewards. Journal of Bone and Mineral Research 22(10), 1492–1501. doi: 10.1359/Jbmr.070518.

Kim, S.W., Lu, Y.H., Williams, E.A., Lai, F., Lee, J.Y., Enishi, T., et al. (2017). Sclerostin Antibody Administration Converts Bone Lining Cells Into Active Osteoblasts. Journal of Bone and Mineral Research 32(5), 892–901. doi: 10.1002/jbmr.3038.

Kim, S.W., Pajevic, P.D., Selig, M., Barry, K.J., Yang, J.Y., Shin, C.S., et al. (2012). Intermittent parathyroid hormone administration converts quiescent lining cells to active osteoblasts. J Bone Miner Res 27(10), 2075–2084. doi: 10.1002/jbmr.1665.

Kujoth, G.C., Hiona, A., Pugh, T.D., Someya, S., Panzer, K., Wohlgemuth, S.E., et al. (2005). Mitochondrial DNA mutations, oxidative stress, and apoptosis in mammalian aging. Science 309(5733), 481–484. doi: 10.1126/science.1112125.

Kuznetsova, A., Brockhoff, P.B., and Christensen, R.H.B. (2017). lmerTest Package: Tests in Linear Mixed Effects Models. Journal of Statistical Software 82(13), 1–26. doi: DOI 10.18637/jss.v082.i13.

Lafuente-Gracia, L., Borgiani, E., Nasello, G., and Geris, L. (2021). Towards in silico Models of the Inflammatory Response in Bone Fracture Healing. Front Bioeng Biotechnol 9, 703725. doi: 10.3389/fbioe.2021.703725.

Lambers, F.M., Schulte, F.A., Kuhn, G., Webster, D.J., and Müller, R. (2011). Mouse tail vertebrae adapt to cyclic mechanical loading by increasing bone formation rate and decreasing bone resorption rate as shown by time-lapsed imaging of dynamic bone morphometry. Bone 49(6), 1340–1350. doi: 10.1016/j.bone.2011.08.035.

Lawson, L.Y., Brodt, M.D., Migotsky, N., Chermside-Scabbo, C.J., Palaniappan, R., and Silva, M.J. (2022). Osteoblast-Specific Wnt Secretion Is Required for Skeletal Homeostasis and Loading-Induced Bone Formation in Adult Mice. Journal of Bone and Mineral Research 37(1), 108–120. doi: 10.1002/jbmr.4445.

Ledoux, C., Boaretti, D., Sachan, A., Muller, R., and Collins, C.J. (2022). Clinical Data for Parametrization of In Silico Bone Models Incorporating Cell-Cytokine Dynamics: A Systematic Review of Literature. Front Bioeng Biotechnol 10, 901720. doi: 10.3389/fbioe.2022.901720.

Levchuk, A., Zwahlen, A., Weigt, C., Lambers, F.M., Badilatti, S.D., Schulte, F.A., et al. (2014). The Clinical Biomechanics Award 2012-Presented by the European Society of Biomechanics: Large scale simulations of trabecular bone adaptation to loading and treatment. Clinical Biomechanics 29(4), 355–362. doi: 10.1016/j.clinbiomech.2013.12.019.

Marques, F.C., Boaretti, D., Walle, M., Scheuren, A.C., Schulte, F.A., and Muller, R. (2023). Mechanostat parameters estimated from time-lapsed in vivo micro-computed tomography data of mechanically driven bone adaptation are logarithmically dependent on loading frequency. Front Bioeng Biotechnol 11, 1140673. doi: 10.3389/fbioe.2023.1140673.

Matsuzaka, T., Matsugaki, A., and Nakano, T. (2021). Control of osteoblast arrangement by osteocyte mechanoresponse through prostaglandin E2 signaling under oscillatory fluid flow stimuli. Biomaterials 279. doi: ARTN 121203 10.1016/j.biomaterials.2021.121203.

McKenzie, J., Smith, C., Karuppaiah, K., Langberg, J., Silva, M.J., and Ornitz, D.M. (2019). Osteocyte Death and Bone Overgrowth in Mice Lacking Fibroblast Growth Factor Receptors 1 and 2 in Mature Osteoblasts and Osteocytes. J Bone Miner Res 34(9), 1660–1675. doi: 10.1002/jbmr.3742.

Meakin, L.B., Galea, G.L., Sugiyama, T., Lanyon, L.E., and Price, J.S. (2014). Age-Related Impairment of Bones’ Adaptive Response to Loading in Mice Is Associated With Sex-Related Deficiencies in Osteoblasts but No Change in Osteocytes. Journal of Bone and Mineral Research 29(8), 1859–1871. doi: 10.1002/jbmr.2222.

Meng, D., Frank, A.R., and Jewell, J.L. (2018). mTOR signaling in stem and progenitor cells. Development 145(1). doi: 10.1242/dev.152595.

Norddahl, G.L., Pronk, C.J., Wahlestedt, M., Sten, G., Nygren, J.M., Ugale, A., et al. (2011). Accumulating mitochondrial DNA mutations drive premature hematopoietic aging phenotypes distinct from physiological stem cell aging. Cell Stem Cell 8(5), 499–510. doi: 10.1016/j.stem.2011.03.009.

Ominsky, M.S., Brown, D.L., Van, G., Cordover, D., Pacheco, E., Frazier, E., et al. (2015). Differential temporal effects of sclerostin antibody and parathyroid hormone on cancellous and cortical bone and quantitative differences in effects on the osteoblast lineage in young intact rats. Bone 81, 380–391. doi: 10.1016/j.bone.2015.08.007.

Ominsky, M.S., Niu, Q.T., Li, C., Li, X., and Ke, H.Z. (2014). Tissue-level mechanisms responsible for the increase in bone formation and bone volume by sclerostin antibody. J Bone Miner Res 29(6), 1424–1430. doi: 10.1002/jbmr.2152.

Pflanz, D., Birkhold, A.I., Albiol, L., Thiele, T., Julien, C., Seliger, A., et al. (2017). deficiency led to a greater cortical bone formation response to mechanical loading and altered gene expression. Scientific Reports 7. doi: ARTN 9435 10.1038/s41598-017-09653-9.

Razi, H., Birkhold, A.I., Weinkamer, R., Duda, G.N., Willie, B.M., and Checa, S. (2015a). Aging Leads to a Dysregulation in Mechanically Driven Bone Formation and Resorption. J Bone Miner Res 30(10), 1864–1873. doi: 10.1002/jbmr.2528.

Razi, H., Birkhold, A.I., Zaslansky, P., Weinkamer, R., Duda, G.N., Willie, B.M., et al. (2015b). Skeletal maturity leads to a reduction in the strain magnitudes induced within the bone: A murine tibia study. Acta Biomaterialia 13, 301–310. doi: 10.1016/j.actbio.2014.11.021.

Scheuren, A.C., D’Hulst, G., Kuhn, G.A., Masschelein, E., Wehrle, E., De Bock, K., et al. (2020a). Hallmarks of frailty and osteosarcopenia in prematurely aged PolgA((D257A/D257A)) mice. J Cachexia Sarcopenia Muscle 11(4), 1121–1140. doi: 10.1002/jcsm.12588.

Scheuren, A.C., Vallaster, P., Kuhn, G.A., Paul, G.R., Malhotra, A., Kameo, Y., et al. (2020b). Mechano-Regulation of Trabecular Bone Adaptation Is Controlled by the Local Environment and Logarithmically Dependent on Loading Frequency. Frontiers in Bioengineering and Biotechnology 8. doi: ARTN 566346 10.3389/fbioe.2020.566346.

Schulte, F.A., Lambers, F.M., Kuhn, G., and Müller, R. (2011). micro-computed tomography allows direct three-dimensional quantification of both bone formation and bone resorption parameters using time-lapsed imaging. Bone 48(3), 433–442. doi: 10.1016/j.bone.2010.10.007.

Schulte, F.A., Ruffoni, D., Lambers, F.M., Christen, D., Webster, D.J., Kuhn, G., et al. (2013a). Local Mechanical Stimuli Regulate Bone Formation and Resorption in Mice at the Tissue Level. Plos One 8(4). doi: ARTN e62172 10.1371/journal.pone.0062172.

Schulte, F.A., Zwahlen, A., Lambers, F.M., Kuhn, G., Ruffoni, D., Betts, D., et al. (2013b). Strain-adaptive in silico modeling of bone adaptation--a computer simulation validated by in vivo micro-computed tomography data. Bone 52(1), 485–492. doi: 10.1016/j.bone.2012.09.008.

Somerville, J.M., Aspden, R.M., Armour, K.E., Armour, K.J., and Reid, D.M. (2004). Growth of C57BL/6 mice and the material and mechanical properties of cortical bone from the tibia. Calcif Tissue Int 74(5), 469–475. doi: 10.1007/s00223-003-0101-x.

Thompson, M.L., Jimenez-Andrade, J.M., and Mantyh, P.W. (2016). Sclerostin Immunoreactivity Increases in Cortical Bone Osteocytes and Decreases in Articular Cartilage Chondrocytes in Aging Mice. J Histochem Cytochem 64(3), 179–189. doi: 10.1369/0022155415626499.

Tourolle, D.C., Dempster, D.W., Ledoux, C., Boaretti, D., Aguilera, M., Saleem, N., et al. (2021). Ten-Year Simulation of the Effects of Denosumab on Bone Remodeling in Human Biopsies. JBMR Plus 5(6), e10494. doi: 10.1002/jbm4.10494.

van Hove, R.P., Nolte, P.A., Vatsa, A., Semeins, C.M., Salmon, P.L., Smit, T.H., et al. (2009). Osteocyte morphology in human tibiae of different bone pathologies with different bone mineral density - Is there a role for mechanosensing? Bone 45(2), 321–329. doi: 10.1016/j.bone.2009.04.238.

Webster, D., Schulte, F.A., Lambers, F.M., Kuhn, G., and Muller, R. (2015). Strain energy density gradients in bone marrow predict osteoblast and osteoclast activity: a finite element study. J Biomech 48(5), 866–874. doi: 10.1016/j.jbiomech.2014.12.009.

Webster, D.J., Morley, P.L., van Lenthe, G.H., and Muller, R. (2008). A novel in vivo mouse model for mechanically stimulated bone adaptation--a combined experimental and computational validation study. Comput Methods Biomech Biomed Engin 11(5), 435–441. doi: 10.1080/10255840802078014.

Xiong, J., Piemontese, M., Onal, M., Campbell, J., Goellner, J.J., Dusevich, V., et al. (2015). Osteocytes, not Osteoblasts or Lining Cells, are the Main Source of the RANKL Required for Osteoclast Formation in Remodeling Bone. PLoS One 10(9), e0138189. doi: 10.1371/journal.pone.0138189.

