## Supplementary Material for "Dysregulated mechanoregulation of load-induced bone adaptation in prematurely aged mice: insights from a micro-multiphysics agent-based model"

Daniele Boaretti *et al.*

**This PDF file includes:**

Supplementary Text
Figs. S1 to S4
Tables S1 to S10
References

**Supplementary Text**

Micro-FE analysis of the *in vivo* and *in silico* images
The bone voxels of the *in vivo* images were identified with a threshold of 580 mg HA/cm$^3$, as performed previously (Scheuren et al., 2020), while the bone voxels of the *in silico* images were identified with the voxels having a mineral concentration of at least 0.5. With such thresholded images, intervertebral discs were digitally added to the proximal and distal ends of these vertebrae. For the definition of the micro-FE models, the discs were modeled as bone voxels. Then, micro-FE models were created by setting the Young's modulus of bone voxels to 14.8 GPa, of marrow voxels to 2 MPa, and of background voxels to 0 MPa. A Poisson ratio of 0.3 was set for all voxels. The application of a compressing force along the longitudinal direction of the sample was simulated. For the loaded groups, the magnitude of the force was set to 8 N, while it was set to 4 N for the control groups, as a physiological compressing force of the CV6 previously estimated (Christen et al., 2012). The nodes at the proximal end of the micro-FE model were set to be fixed, while the nodes at the distal ends were set to have a 1% displacement along the direction of the simulated force. The micro-FE analysis was carried out on these images to compute the SED using ParOsol (Flaig and Arbenz, 2011). The Swiss National Supercomputing Center (CSCS, Lugano, Switzerland) was used for running the analysis. Each micro-FE analysis took less than a minute, employing two nodes (Piz Daint, a Cray XC40/50 model).

Extension of the in silico micro-MPA model for simulating mechanical loading and aging
The original version of the *in silico* micro-MPA model was used for studying trabecular bone remodeling in the sixth caudal vertebra in a physiological condition, with the application of a compressing force of 4 N to vertebra along the longitudinal direction (Boaretti et al., 2023). The magnitude of this force was defined previously (Christen et al., 2012), who estimated it in the same mechanical environment.
The computational analysis of the mechanical environment was performed following an established procedure briefly mentioned in the Methods section. From the micro-FE analysis, the effective strain in the trabecular region was gaussian dilated for the bone voxels ($\varepsilon_{bone}$) as well as for the marrow voxels ($\varepsilon_{marrow}$) with a sigma of 2.5. These mechanical properties were used for the definition of the mechanomic profiles and for converting the mechanical input into the production of cytokines according to a catabolic or anabolic response.
When simulating mechanical loading, the micro-FE models were applied twice for the definition of the mechanical environment, to simulate the actual application of the mechanical loading in the experimental setup. Therefore, at the beginning of the analysis, the same micro-FE model was used to determine the mechanical environment with the physiological compressing force of 4 N prior to loading and with the force of 8 N when the loading study actually starts.
The single-cell mechanomics profiles were rescaled to take into account bone growth and adaptation to a higher range of effective strains at the end of the loading period. The definition of the mechanostat threshold for the single-cell mechanomics of the osteocytes in the physiological condition, i.e. when the simulated force had a magnitude of 4N, was defined as follows: $mech_{thres}^{bone} = 0.95 * median(\varepsilon_{bone})$, where $\varepsilon_{bone}$ is the local mechanical signal in the trabecular region of the given vertebra (Boaretti et al., 2023). In the new conditions we simulated in this

study, the application of the mechanical force in the loading condition was introduced in the single-cell mechanomics of the osteocytes as follows: $mech_{loading}^{bone} = mech_{thres}^{bone} \times \frac{applied\_force}{physiological\_force}$, where $physiological\_force$ is 4 N, $applied\_force$ is 8 N which corresponds to the peak load used in the experimental analysis.
The definition of the mechanostat threshold for the single-cell mechanomics of the osteoblasts and osteoclasts in the physiological condition was defined as follows: $mech_{thres}^{marrow} = 0.45 * median(\varepsilon_{marrow})$, where $\varepsilon_{marrow}$ is the local mechanical signal in the trabecular region of the marrow of the given vertebra. The application of the mechanical force in the loading condition was introduced in the formulation of the single-cell mechanomics of the osteoblasts and osteoclasts as follows: $mech_{loading}^{marrow} = mech_{thres}^{marrow} \times \frac{applied_{force}}{physiological\_force}$.
Taking into account a possible aging and loading effect on the single-cell mechanomics of the cells, we introduced a variable $\alpha_{aging}^{loading}$ >1 to increase the single-cell mechanomics thresholds defined above for the osteocytes, leading to the following formula: $mech_{loading,aging}^{bone} = mech_{loading}^{bone} \times \alpha_{aging}^{loading}$.
The simulations followed the established model generation and initial execution to reach a stable remodeling phase as described previously (Boaretti et al., 2023). This step was carried out with the corresponding control configuration for the sample's age. After this phase, the configuration was updated using the corresponding parameters of the model, loading force, and single-cell mechanomics described above.

Configurations for the four branches

The four groups were simulated primarily by changing the activity of the cells. The changes in the single-cell production of osteoprotegerin (OPG), receptor activator of nuclear factor kB ligand (RANKL) and sclerostin (Scl) by osteoblasts (Ob), lining cells, and osteocytes (Ot) were required due to the different cell densities and cell availability rates modelled here. We manipulated the apoptosis, proliferation, and bone formation-related parameters. The full list of values is reported in Supplementary Table S4. The initial non-surface cell (MSC, HSC) density values for the control configurations were $1.25 \times 10^7$ $1/ml$ and $8.75 \times 10^6$ $1/ml$ for the young and old mice, respectively.

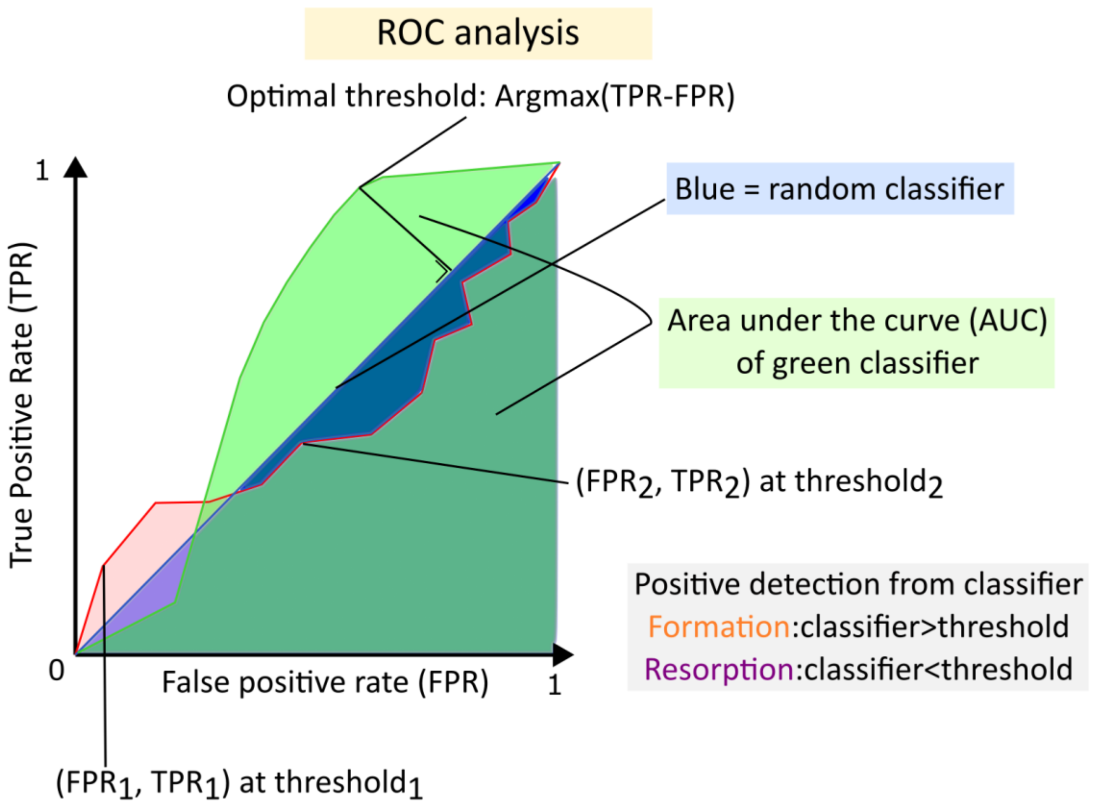


**Fig. S1.**

**The ROC-AUC analysis and the optimal threshold computation.** Given a classifier of binary events, it is possible to obtain ROC curve by changing the threshold of the mechanical signal for classifying the events: on each voxel of the trabecular surface, bone formation events were predicted when the mechanical signal value was greater than a threshold whereas bone resorption events were predicted when the mechanical signal value was lower than a threshold. As a consequence, for each threshold a true positive rate (TPR) and a false positive rate (FPR) are obtained, and these pairs can be plotted to obtain the ROC curve. The area under the ROC curve (AUC) provides an estimate of the quality of the classifier. The optimal threshold of the classifier was obtained by maximizing the difference between TPR and FPR.

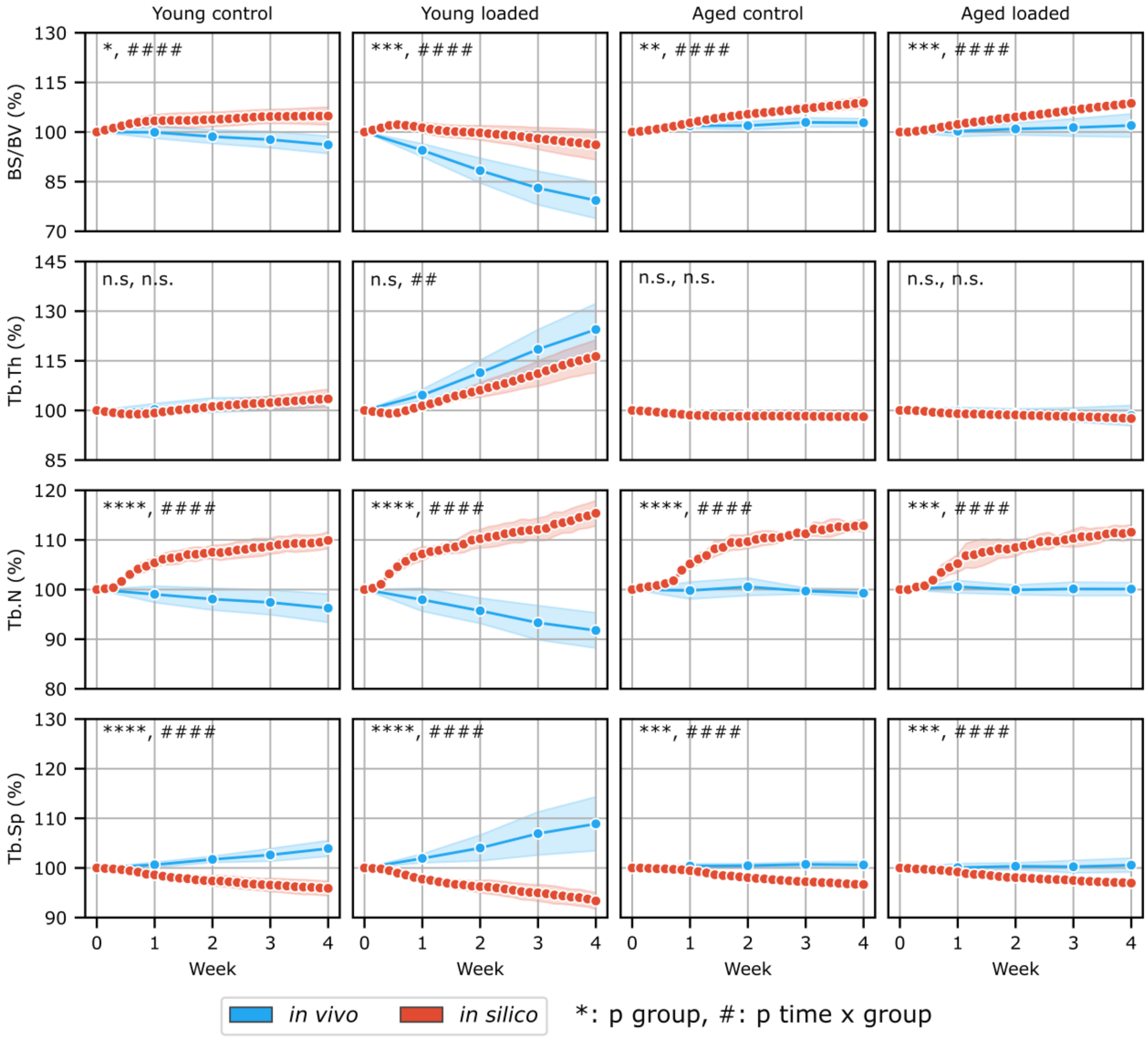


**Fig. S2.**

Normalized specific bone surface (Norm. BS/BV), trabecular thickness (Norm. Tb.Th), trabecular number (Norm. Tb.N) and trabecular spacing (Norm. Tb.Sp) in the four groups of mice *in silico* and *in vivo* data over four weeks. The *in vivo* data show the data from the time-lapsed weekly images while the *in silico* data are reported on a daily basis. Light blue = *in vivo*, Red = *in silico*. n=5 *in silico* and *in vivo* control groups, n=9 *in vivo* loaded groups. (n.s., * $p<0.05$, ** $p<0.01$, *** $p<0.001$, **** $p<0.0001$ for the group comparison, # $p<0.05$, ## $p<0.01$, ### $p<0.001$, #### $p<0.0001$ for the interaction time*group determined by two-way ANOVA).

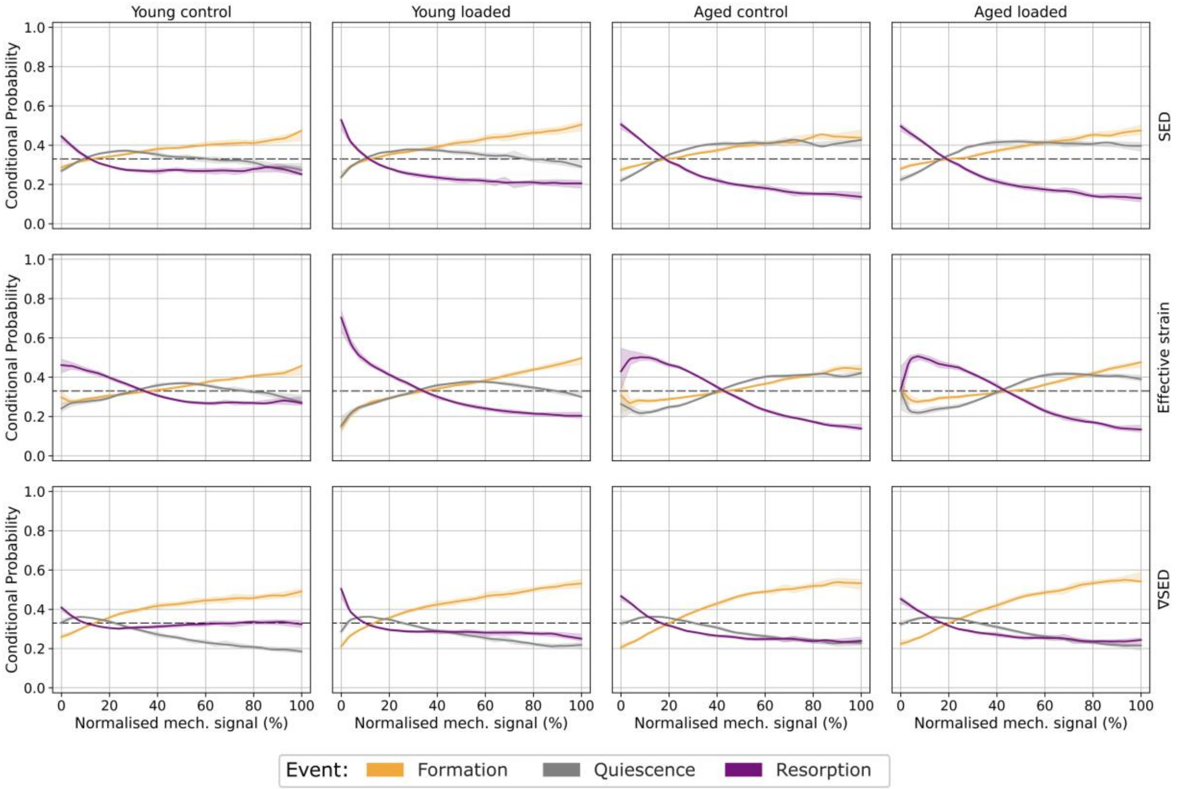


**Fig. S3.**

The mechanoregulation analysis in the trabecular region of the sixth caudal vertebra for the four groups of PolgA mice *in vivo*. The first, second, and third rows show the mechanoregulation curves of formation, resorption and quiescence using strain energy density values in bone ($SED_b$), effective strain in bone ($EFF_b$), and the norm of the gradient of strain energy density ($\nabla SED_m$), respectively, for the first weekly interval. The orange, gray and purple curves are related to bone formation, quiescence and resorption respectively. The dashed line represents the random probability of observing one of these three events, corresponding to one third.

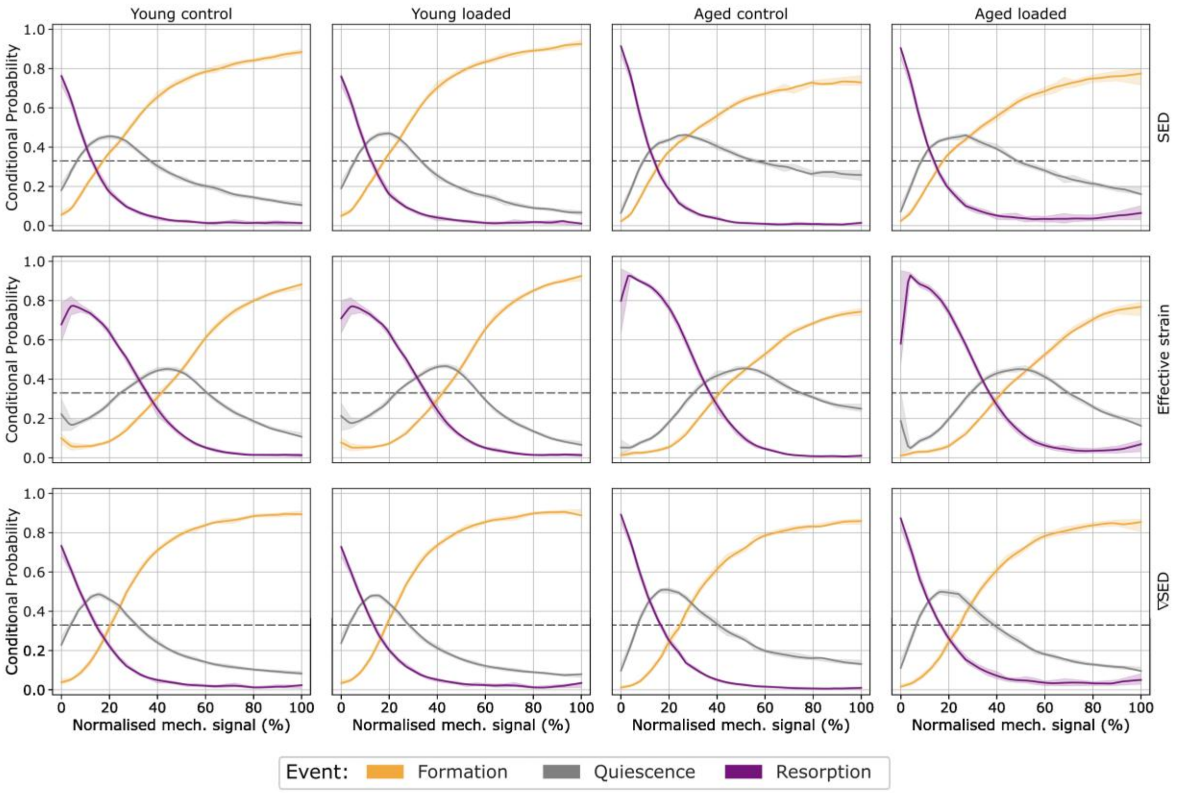


**Fig. S4.**

The mechanoregulation analysis in the trabecular region of the sixth caudal vertebra for the four groups of PolgA mice *in silico*. The first, second, and third rows show the mechanoregulation curves of formation, resorption and quiescence using strain energy density values in bone ($SED_b$), effective strain in bone ($EFF_b$), and the norm of the gradient of strain energy density ($\nabla SED_m$), respectively, for the first weekly interval. The orange, gray and purple curves are related to bone formation, quiescence and resorption respectively. The dashed line represents the random probability of observing one of these three events, corresponding to one third.

**Table S1.**

| Parameter | Weeks | Young control | | Young loaded | |
|---|---|---|---|---|---|
| | | *In vivo* | *In silico* | *In vivo* | *In silico* |
| BFR (%/day) | 0-2 | 0.751±0.266 | 0.596±0.066 | 1.251±0.317 | 1.048±0.080 |
| | 2-4 | 0.693±0.195 | 0.344±0.019 | 1.006±0.263 | 1.132±0.268 |
| | p-val | n.s., n.s. | | n.s., n.s. | |
| BRR (%/day) | 0-2 | 1.004±0.250 | 0.748±0.379 | 0.721±0.276 | 0.851±0.345 |
| | 2-4 | 0.760±0.220 | 0.470±0.135 | 0.698±0.213 | 0.761±0.155 |
| | p-val | p<0.01, n.s. | | n.s., n.s. | |
| MAR (μm/day) | 0-2 | 1.034±0.092 | 1.075±0.004 | 1.179±0.177 | 1.308±0.031 |
| | 2-4 | 0.972±0.077 | 0.839±0.021 | 1.049±0.205 | 1.187±0.028 |
| | p-val | n.s., n.s. | | p<0.05, n.s. | |
| MRR (μm/day) | 0-2 | 1.908±0.157 | 1.949±0.018 | 1.941±0.270 | 1.953±0.042 |
| | 2-4 | 1.709±0.037 | 1.583±0.136 | 1.945±0.152 | 1.742±0.029 |
| | p-val | n.s., p<0.05 | | n.s., p<0.01 | |
| MS (%) | 0-2 | 46.460±4.625 | 28.712±1.491 | 57.744±4.724 | 37.798±0.373 |
| | 2-4 | 45.501±3.219 | 26.625±1.990 | 59.567±4.952 | 44.341±0.722 |
| | p-val | p<0.0001, n.s. | | p<0.01, n.s. | |
| ES (%) | 0-2 | 28.318±4.757 | 19.923±4.367 | 20.129±4.696 | 20.481±4.794 |
| | 2-4 | 27.866±1.661 | 15.573±3.613 | 22.030±3.776 | 22.393±2.614 |
| | p-val | p<0.0001, n.s. | | n.s., n.s. | |

Median and interquartile range of the dynamic bone morphometry for the in silico and in vivo young groups with a 2-week interval. P-values from the group and the interaction time-group two-way ANOVA are also reported.

**Table S2.**

| Parameter | Weeks | Aged control | | Aged loaded | |
|---|---|---|---|---|---|
| | | *In vivo* | *In silico* | *In vivo* | *In silico* |
| BFR (%/day) | 0-2 | 0.188±0.035 | 0.139±0.020 | 0.183±0.053 | 0.135±0.040 |
| | 2-4 | 0.204±0.112 | 0.142±0.020 | 0.188±0.124 | 0.066±0.040 |
| | p-val | p<0.05, n.s. | | n.s., p<0.05 | |
| BRR (%/day) | 0-2 | 0.306±0.087 | 0.241±0.044 | 0.165±0.152 | 0.237±0.031 |
| | 2-4 | 0.247±0.133 | 0.214±0.037 | 0.295±0.195 | 0.204±0.012 |
| | p-val | n.s., n.s. | | n.s., n.s. | |
| MAR (μm/day) | 0-2 | 0.831±0.085 | 0.848±0.003 | 0.777±0.031 | 0.942±0.005 |
| | 2-4 | 0.785±0.096 | 0.789±0.007 | 0.802±0.102 | 0.794±0.027 |
| | p-val | n.s., p<0.05 | | p<0.001, p<0.0001 | |
| MRR (μm/day) | 0-2 | 0.901±0.095 | 1.217±0.010 | 0.799±0.063 | 1.130±0.011 |
| | 2-4 | 0.821±0.104 | 1.062±0.033 | 0.878±0.153 | 1.010±0.006 |
| | p-val | p<0.01, p<0.0001 | | p<0.001, p<0.0001 | |
| MS (%) | 0-2 | 30.562±1.000 | 19.661±3.530 | 30.898±3.584 | 17.677±1.001 |
| | 2-4 | 31.325±2.355 | 16.512±2.949 | 31.262±8.838 | 9.030±3.130 |
| | p-val | p<0.001, p<0.01 | | p<0.001, p<0.01 | |
| ES (%) | 0-2 | 38.869±4.312 | 18.428±1.452 | 32.536±4.052 | 18.050±0.400 |
| | 2-4 | 34.021±5.209 | 18.586±1.793 | 32.932±9.474 | 18.395±0.811 |
| | p-val | p<0.001, n.s. | | p<0.0001, n.s. | |

Median and interquartile range of the dynamic bone morphometry for the in silico and in vivo aged groups with a 2-week interval. P-values from the group and the interaction time-group two-way ANOVA are also reported..

**Table S3.**

| | Young control | | Young loaded | |
|---|---|---|---|---|
| | **Group** | **Time x group** | **Group** | **Time x group** |
| Norm. BV/TV | n.s. | n.s. | n.s. | n.s. |
| Norm. BS/BV | p<0.05 | p<0.0001 | p<0.01 | p<0.0001 |
| Norm. Tb.Th | n.s. | n.s. | n.s. | p<0.01 |
| Norm. Tb.Sp | p<0.0001 | p<0.0001 | p<0.0001 | p<0.0001 |
| Norm. Tb.N | p<0.0001 | p<0.0001 | p<0.0001 | p<0.0001 |
| BFR (%/day) | n.s. | n.s. | n.s. | n.s. |
| BRR (%/day) | p<0.01 | n.s. | n.s. | n.s. |
| MAR (μm/day) | n.s. | n.s. | p<0.05 | n.s. |
| MRR (μm/day) | n.s. | p<0.05 | n.s. | p<0.01 |
| MS (%) | p<0.0001 | n.s. | p<0.01 | n.s. |
| ES (%) | p<0.0001 | n.s. | n.s. | n.s. |

Complete table of the p-values related to the statistical difference between *in vivo* and *in silico* static and dynamic bone morphometric parameters of the young groups of mice, showing the comparisons between groups («Group» columns) and for the interaction time-group («Time x group» columns).

**Table S4.**

| | Aged control | | Aged loaded | |
|---|---|---|---|---|
| | **Group** | **Time x group** | **Group** | **Time x group** |
| Norm. BV/TV | n.s. | n.s. | n.s. | n.s. |
| Norm. BS/BV | p<0.01 | p<0.0001 | p<0.001 | p<0.0001 |
| Norm. Tb.Th | n.s. | n.s. | n.s. | n.s. |
| Norm. Tb.Sp | p<0.001 | p<0.0001 | p<0.001 | p<0.0001 |
| Norm. Tb.N | p<0.0001 | p<0.0001 | p<0.001 | p<0.0001 |
| BFR (%/day) | p<0.05 | n.s. | n.s. | p<0.05 |
| BRR (%/day) | n.s. | n.s. | n.s. | n.s. |
| MAR (μm/day) | n.s. | p<0.05 | p<0.001 | p<0.0001 |
| MRR (μm/day) | p<0.01 | p<0.001 | p<0.001 | p<0.001 |
| MS (%) | p<0.001 | p<0.01 | p<0.001 | p<0.01 |
| ES (%) | p<0.001 | n.s. | p<0.0001 | n.s. |

Complete table of the p-values related to the statistical difference between *in vivo* and *in silico* static and dynamic bone morphometric parameters of the aged groups of mice, showing the comparisons between groups («Group» columns) and for the interaction time-group («Time x group» columns).

**Table S5.**

| Linear regression line: Norm. Tb.Th (%) = m·x + q | | | | | |
|---|---|---|---|---|---|
| **x** | **Group** | **$R^2$** | **Slope (m)** | **Intercept (q)** | **p value** |
| Ob.N/Oc.N(-) | young control | 0.06 | 0.0389 | 99.9 | p<0.01 |
| | young loaded | 0.27 | -0.3014 | 114.4 | p<0.0001 |
| | aged control | 0.58 | 0.5025 | 95.8 | p<0.0001 |
| | aged loaded | 0.61 | 0.4551 | 96.9 | p<0.0001 |
| Oc.N/BS (cells/$mm^2$) | young control | 0.04 | -0.0063 | 102.1 | p<0.05 |
| | young loaded | 0.82 | 0.0419 | 93.8 | p<0.0001 |
| | aged control | 0.54 | -0.0082 | 103.2 | p<0.0001 |
| | aged loaded | 0.57 | -0.0072 | 103.2 | p<0.0001 |
| Scl (pg/ml) | young control | 0.23 | 0.1346 | 93.3 | p<0.0001 |
| | young loaded | 0.51 | 0.4222 | 80.8 | p<0.0001 |
| | aged control | 0.01 | -0.0120 | 99.5 | n.s. |
| | aged loaded | 0.18 | -0.0541 | 103.5 | p<0.0001 |

Linear regression results between selected output variables of the micro-MPA model and normalized trabecular thickness (Norm. Tb.Th).

**Table S6.**

| Group | Variable | Paired t-test p values | Cohen's D |
|---|---|---|---|
| Young control | AUC formation | $p<0.001$ | 5.79 |
| | AUC resorption | $p<0.0001$ | 4.34 |
| Young loaded | AUC formation | $p<0.0001$ | 5.43 |
| | AUC resorption | $p<0.0001$ | 5.75 |
| Aged control | AUC formation | $p<0.0001$ | 5.14 |
| | AUC resorption | $p<0.0001$ | 6.72 |
| Aged loaded | AUC formation | $p<0.0001$ | 5.57 |
| | AUC resorption | $p<0.0001$ | 6.39 |

Statistical difference between the area under the curve (AUC) for formation and resorption computed with the strain energy density in bone **(**$SED_b$) and the norm of the gradient of the SED in the marrow ($\nabla SED_m$) from paired t-test and Cohen's D.

**Table S7.**

| | Young control | | | Young loaded | | |
|---|---|---|---|---|---|---|
| | **Group** | **Time** | **Time x group** | **Group** | **Time** | **Time x group** |
| $SED_b$ | n.s. | n.s. | n.s. | n.s. | $p<0.0001$ | n.s. |
| $\nabla SED_m$ | n.s. | n.s. | n.s. | n.s. | $p<0.0001$ | n.s. |

Statistical difference between the normalized optimal threshold of formation (Norm.OTF) and optimal threshold of resorption (Norm.OTR) using the strain energy density in bone **(**$SED_b$) and the norm of the gradient of the SED in the marrow ($\nabla SED_m$) in the young groups.

**Table S8.**

| | Aged control | | | Aged loaded | | |
|---|---|---|---|---|---|---|
| | **Group** | **Time** | **Time x group** | **Group** | **Time** | **Time x group** |
| $SED_b$ | n.s. | n.s | n.s. | n.s. | n.s. | n.s. |
| $\nabla SED_m$ | n.s. | n.s | n.s. | n.s. | n.s. | n.s. |

Statistical difference between the normalized optimal threshold of formation (OTF) and optimal threshold of resorption (OTR) using the strain energy density in bone **(**$SED_b$) and the norm of the gradient of the SED in the marrow ($\nabla SED_m$) in the aged groups.

**Table S9.**

| | | Formation | | Resorption | | Net bone response | |
|---|---|---|---|---|---|---|---|
| **Regression line** | | F= y0+a*(1-exp(-b*MS/$MS_{max}$) | | R= y0+a*(exp(-b*MS/$MS_{max}$) | | N= y0+a*(1-exp(-b*MS/$MS_{max}$) | |
| **Mechanical signal (MS)** | | SED | SED gradient | SED | SED gradient | SED | SED gradient |
| **Young control** | y0 | 29.788 | 21.988 | 27.030 | 29.597 | -14.171 | -18.197 |
| | a | 19.462 | 32.418 | 18.388 | 12.577 | 29.253 | 40.970 |
| | b | 0.013 | 0.033 | 0.107 | 0.150 | 0.050 | 0.050 |
| | $R^2$ | 0.917 | 0.978 | 0.885 | 0.800 | 0.915 | 0.956 |
| **Young loaded** | y0 | 28.197 | 21.363 | 21.080 | 25.212 | -18.400 | -20.793 |
| | a | 29.091 | 39.920 | 27.660 | 19.924 | 46.009 | 54.623 |
| | b | 0.012 | 0.0254 | 0.071 | 0.092 | 0.038 | 0.037 |
| | $R^2$ | 0.960 | 0.978 | 0.952 | 0.906 | 0.950 | 0.959 |
| **Aged control** | y0 | 27.639 | 19.132 | 13.254 | 23.878 | -23.214 | -26.579 |
| | a | 24.100 | 38.580 | 38.127 | 22.907 | 57.503 | 58.606 |
| | b | 0.013 | 0.024 | 0.037 | 0.055 | 0.028 | 0.033 |
| | $R^2$ | 0.940 | 0.976 | 0.993 | 0.976 | 0.986 | 0.980 |
| **Aged loaded** | y0 | 28.453 | 20.959 | 11.931 | 23.652 | -21.401 | -23.040 |
| | a | 57.663 | 42.924 | 38.841 | 21.495 | 62.071 | 59.313 |
| | b | 0.004 | 0.017 | 0.034 | 0.049 | 0.022 | 0.026 |
| | $R^2$ | 0.961 | 0.986 | 0.992 | 0.975 | 0.989 | 0.984 |

Summary of non-linear regression functions and corresponding coefficients for the conditional probability (SED in bone and ∇SED in marrow) in trabecular bone for the four groups for the first weekly analysis.

**Table S10.**

| | Young | | Aged | |
|---|---|---|---|---|
| | **Control** | **Loaded** | **Control** | **Loaded** |
| $\beta_{Ob}^{osteoid}$ | $0.100\ zmol/day$ | $0.160\ zmol/day$ | $0.084\ zmol/day$ | $0.090\ zmol/day$ |
| $\beta_{Ob,Li.Ce}^{RANKL}$ | $4.000\ \times\ 10^{-3}\ zmol/day$ | | $6.000 \times 10^{-3}\ zmol/day$ | |
| $\beta_{Ob,Li.Ce}^{OPG}$ | $2.660\ \times\ 10^{-4}\ zmol/day$ | | $4.000\ \times 10^{-4}\ zmol/day$ | |
| $\beta_{Ot}^{Scl}$ | $6.711\ \times\ 10^{-3}\ zmol/day$ | | $1.200\ \times 10^{-2}\ zmol/day$ | |
| $\beta_{Ot}^{RANKL}$ | $1.677 \times\ 10^{-2}\ zmol/day$ | | $1.000\ \times 10^{-2}\ zmol/day$ | |
| $\beta_{Ot}^{OPG}$ | $2.013 \times\ 10^{-2}\ zmol/day$ | | $1.200\ \times 10^{-2}\ zmol/day$ | |
| $\beta_{Oc}^{mineral,osteoid}$ | $1.120\ zmol/day$ | | $0.200\ zmol/day$ | $0.175\ zmol/day$ |
| $\sigma_{Oc}$ | 0.65 | | 0.65 | 0.25 |
| $s^{\mathrm{TGF-\beta1}}$ | 1.2 | 2 | 1 | |
| $s^{\mathrm{RANK}}$ | 1.2 | 1 | | |
| $A_{Oc}$ | $0.005\ 1/day$ | | $0.003\ 1/day$ | |
| $A_{Ob}$ | $0.010\ 1/day$ | | $0.013\ 1/day$ | |
| $P_{MSC}$ | $0.064\ 1/day$ | $0.080\ 1/day$ | $0.080\ 1/day$ | |
| $P_{HSC}$ | $0.064\ 1/day$ | $0.080\ 1/day$ | | |
| $P_{Ob}$ | $0.010\ 1/day$ | $0.0275\ 1/day$ | $0.009\ 1/day$ | |
| $r^{mineral}$ | $2.5\ 1/day$ | | $1.0\ 1/day$ | $2.5\ 1/day$ |
| $\alpha_{aging}^{loading}$ | 1 | | 1.10000 | 1.19333 |

Parameter values for the four different simulation branches.

References


Boaretti, D., Marques, F.C., Ledoux, C., Singh, A., Kendall, J.J., Wehrle, E., et al. (2023). Trabecular bone remodeling in the aging mouse: A micro-multiphysics agent-based model using single-cell mechanomics. *Frontiers in Bioengineering and Biotechnology* 11. doi: ARTN 109129410.3389/fbioe.2023.1091294.

Christen, P., van Rietbergen, B., Lambers, F.M., Muller, R., and Ito, K. (2012). Bone morphology allows estimation of loading history in a murine model of bone adaptation. *Biomech Model Mechanobiol* 11(3-4)**,** 483-492. doi: 10.1007/s10237-011-0327-x.

Flaig, C., and Arbenz, P. (2011). A scalable memory efficient multigrid solver for micro-finite element analyses based on CT images. *Parallel Computing* 37(12)**,** 846-854. doi: 10.1016/j.parco.2011.08.001.

Scheuren, A.C., D'Hulst, G., Kuhn, G.A., Masschelein, E., Wehrle, E., De Bock, K., et al. (2020). Hallmarks of frailty and osteosarcopenia in prematurely aged PolgA((D257A/D257A)) mice. *J Cachexia Sarcopenia Muscle* 11(4)**,** 1121-1140. doi: 10.1002/jcsm.12588.
